# Extended Graphical Lasso for Multiple Interaction Networks for High Dimensional Omics Data

**DOI:** 10.1101/2021.02.16.431400

**Authors:** Yang Xu, Hongmei Jiang, Wenxin Jiang

## Abstract

There has been a spate of interest in association networks in biological and medical research, for example, genetic interaction networks. In this paper, we propose a novel method, the extended joint hub graphical lasso (EDOHA), to estimate multiple related interaction networks for high dimensional omics data across multiple distinct classes. To be specific, we construct a convex penalized log likelihood optimization problem and solve it with an alternating direction method of multipliers (ADMM) algorithm. The proposed method can also be adapted to estimate interaction networks for high dimensional compositional data such as microbial interaction networks. The performance of the proposed method in the simulated studies shows that EDOHA has remarkable advantages in recognizing class-specific hubs than the existing comparable methods. We also present three applications of real datasets. Biological interpretations of our results confirm those of previous studies and offer a more comprehensive understanding of the underlying mechanism in disease.

**Author summary:** Reconstruction of multiple association networks from high dimensional omics data is an important topic, especially in biology. Previous studies focused on estimating different networks and detecting common hubs among all classes. Integration of information over different classes of data while allowing difference in the hub nodes is also biologically plausible. Therefore, we propose a method, EDOHA, to jointly construct multiple interaction networks with capacity in finding different hub networks for each class of data. Simulation studies show the better performance over conventional methods. The method has been demonstrated in three real world data.

## Introduction

With advances in high-throughput sequencing and omics technologies, biological information is being collected at an amazing rate, which stimulates researchers to discover modular structure, relationships and regularities in complex data. Interactions between various biological nodes (e.g. genes, proteins, metabolites) on different levels (e.g. gene regulation, cell signalling) can be represented as graphs and, thus, analysis of such networks might shed new light on the function of biological systems. Hubs, the highly connected nodes at the tail of the power law degree distribution, are known to play a crucial role in biological networks. Some studies have shown that scale-free topology exists in many different organizational levels, such as metabolic networks [1] and cellular networks [2]. Hub nodes may be the most essential elements for community stability and play an important role in the infection and pathogenesis of the virus.

The objective of our research is to estimate multiple interaction networks for high dimensional omics data (e.g. genomics, metagenomics, proteomics and metabolomics) across multiple classes. A common characteristic of the omics data is the deficiency of independent samples (*n*) in comparison with the abundance of features (*p*), that is to say, *p* ≫ *n*. There have been a number of studies proposed to construct interaction networks in the high dimensional setting. Meinshausen and Buhlmann [3] present neighborhood selection to discover network structures. Friedman et al. [4] propose the graphical lasso algorithm to estimate networks using the LASSO penalty. Fan et al. [5] introduce nonconcave penalties and the adaptive LASSO penalty to explore networks. Nevertheless, aforementioned methods are used to depict the relationship networks between features for one class only. When there are multiple classes, such as healthy and diseased conditions, a straightforward method is to construct the network for each class separately and then compare their differences. However, these procedures may sacrifice the similarity shared between multiple classes, which may be critically important to find out the principal elements related to the disease. One would expect these networks to be similar to each other, since they are from the same type of entities. The joint graphical lasso (JGL) [6] is proposed to estimate multiple models simultaneously, which ignores the scale-free network and is unable to detect hubs explicitly. In the model of JRmGRN [7], it identifies common hub elements across multiple classes by jointly using distinct datasets. In many situations, hub nodes that are specific to an individual network also exist. For example, in the tissue-specific networks associated with SARS-CoV-2, both common and class-specific key hubs are revealed in diverse tissues [8]. Common hub features are essential to all class and class-specific hubs could convey particular biology information. This inspires us to explore a new model to incorporate both common and class-specific hub nodes when jointly constructing interaction networks.

The proposed method can be applied to any omics data which follow multivariate normal distribution. It can also be easily adapted to study multiple interaction networks for high dimensional compositional data such as microbial networks by employing some suitable transformation. The performance of the proposed method and comparison with other methods will be evaluated by simulation studies for compositional data and real data analysis.

## Materials and methods

Gaussian graphical models (GGMs) are now frequently used to describe biological feature association networks and to detect conditionally dependent features. Correlation networks could be expressed as an undirected, weighted graph *ℊ* = (**V, E**) where the vertex set **V** = {*v*_1_, *v*_2_, … , *v*_*p*_}represents the *p* feature nodes (e.g., genes, microbes or proteins) and the edge set **E** contains the possible associations among nodes. Suppose the observations (suitably transformed if necessary) (*r*_1_, … , *r*_*p*_) are drawn from a multivariate normal distribution with covariance **Σ**, the non-zero elements of the off-diagonal entries of the inverse covariance matrix **Θ** = **Σ**^*−*1^ define the adjacency matrix of the graph *ℊ* and thus describe the factorization of the normal distribution into conditionally dependent components [9]. Because the number of samples *n* is smaller than the number of features *p* and **Θ** is expected to be sparse, penalized maximum likelihood approaches are proposed to estimate the precision matrix **Σ**^*−*1^, which yields a sparse estimation of precision matrix 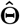.

### The general formulation for EDOHA

We present the extended joint hub graphical lasso (EDOHA) algorithm for constructing multiple interaction networks from multiple classes. Suppose that there are *K* classes of data sets, corresponding to *K* different levels of a phenotype variable or *K* different conditions, such as control group, carrier group and disease group. Let 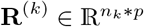 be a matrix representing the data of *p* features and *n*_*k*_ samples for *k*th class. Assume that the observations (suitably transformed if necessary) are independent, identically distributed: 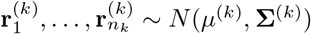, where **r**^(*k*)^ represents biological data from the *k*th class. The log likelihood for the data takes the form

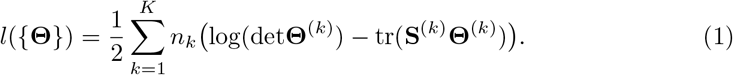

where **S**^(*k*)^ is the empirical covariance estimation of **r**^(*k*)^. The non-zero element 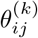 in 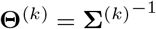 indicates node *i* and *j* for the *k*th class are conditionally dependent. Most elements in **Θ**^(*k*)^ are expected to be zero. JRmGRN [7] has decomposed the precision matrix **Θ**^(*k*)^ into two parts: the elementary symmetric network for the *k*th class **Z**^(*k*)^, mainly containing the non-hub node correlation information, and the network for hub nodes **V**, where **V** is a matrix with entirely zero or almost completely nonzero columns, so that a few hub nodes are expected to have a large number of interactions with many other nodes. Considering that some of the hub codes are common among all classes and others are specific to different classes, we replace the same network **V** with **V**^(*k*)^ for the *k*th class, including common and class-specific hub correlation information. Our method aims to investigate these class-specific hub nodes explicitly. To estimate {**Θ**} = (**Θ**^(1)^, **Θ**^(2)^, … , **Θ**^(*K*)^) when *p > n*_*k*_, we take a penalized log likelihood approach

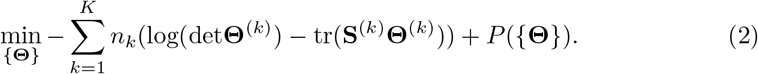

The penalty function *P* ({**Θ**}) has the following form,

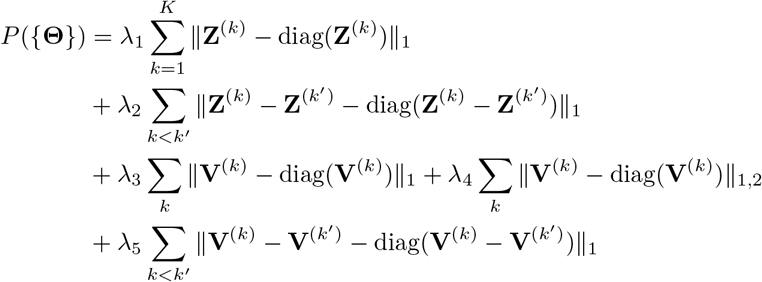

where **Z**^(*k*)^ + **V**^(*k*)^ + (**V**^(*k*)^)^*T*^ = **Θ**^(*k*)^, and 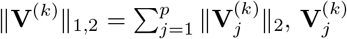 is the *j*th column of matrix **V**^(*k*)^. Here *λ*_1_, *λ*_2_, *λ*_3_, *λ*_4_, *λ*_5_ are five nonnegative tuning parameters. *λ*_1_ and *λ*_3_ control the sparsity of elementary network **Z**^(*k*)^ and hub network **V**^(*k*)^ respectively. *λ*_4_ allows **V**^(*k*)^ to have zero columns and dense non-zero columns, where the non-zero columns represent the respective hub nodes in *k*th class. And *λ*_2_, *λ*_5_ encourage the elementary networks and hub networks to have the similarity. When *λ*_1_, *λ*_2_, *λ*_3_, *λ*_4_ and *λ*_5_ are fixed, the expression of (2) is a convex optimization problem, which can be solved by efficient algorithms. The convexity of (2) is based on the following facts: both negative log determinant and norm functions are convex functions, so is the nonnegative combination of convex functions.

#### Remark 1.

*JRmGRN has four parameters, which accommodate connectivity levels among non-hubs in each class, similarity between non-hubs networks, different numbers of hubs and sparsity levels of hubs. It decomposes the precision matrix into elementary network unique to each class and common hub network, which is equipped with the ability to identify common hubs. Its penalty function is*

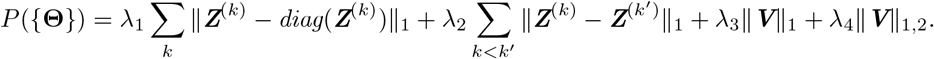

*Compared with the JRmGRN model, EDOHA replaces the common hub network with respective hub network for each class and thus we are able to find out the common and class-specific hub nodes simultaneously. It is easy to find that JRmGRN is a sub-case of EDOHA when λ*_5_ *is large enough. Common hub features across multiple classes could be crucial to regulate biological interaction, while class-specific hubs may mediate specific phenotype. Our proposed method may help to explain which features play a significant part in different phenotypic traits or in different conditions*.

### An ADMM algorithm for EDOHA

We solve the problem using an alternating directions method of multipliers algorithm [10], which allows us to decouple some of the terms that are difficult to optimize jointly. We assume that **Θ**^(*k*)^ is positive definite for *k* = 1, … , *K*. We note that the problem can be reformulated as a consensus problem [11]:

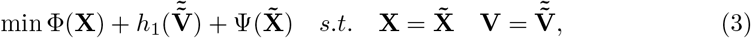

where X=(**Θ**^(1)^, **Z**^(1)^, **V**^(1)^, **…**,**Θ**^(*k*)^, **Z**^(*k*)^, **V**^(*k*)^), 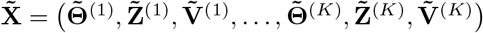, and

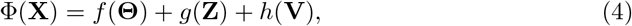

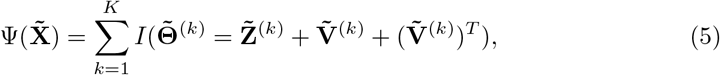

where

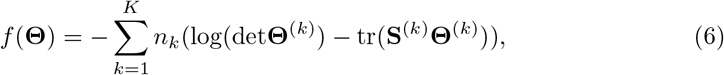

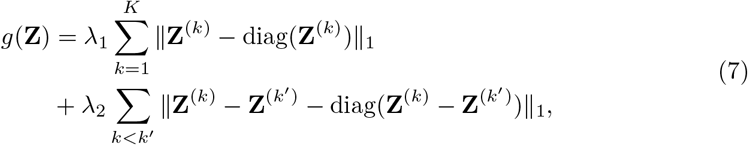

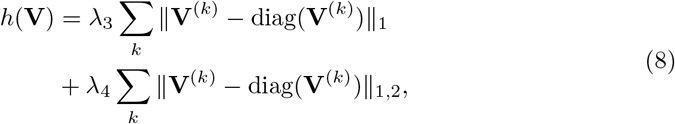

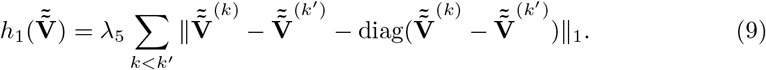

And

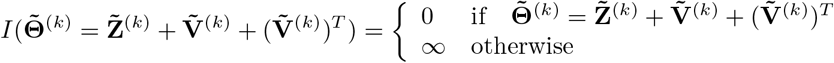

The scaled augmented Lagrangian is given by

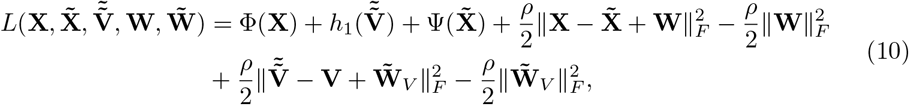

where **X**, 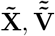 are the primal variables, 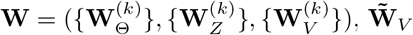 are the dual variables. 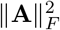 denotes the Frobenius norm of **A**. Here *ρ* is a positive parameter for the scaled Lagrangian form. We set *ρ* = 2.5 as is used in Deng et al. [7].

The iteration of ADMM can be described as follows:

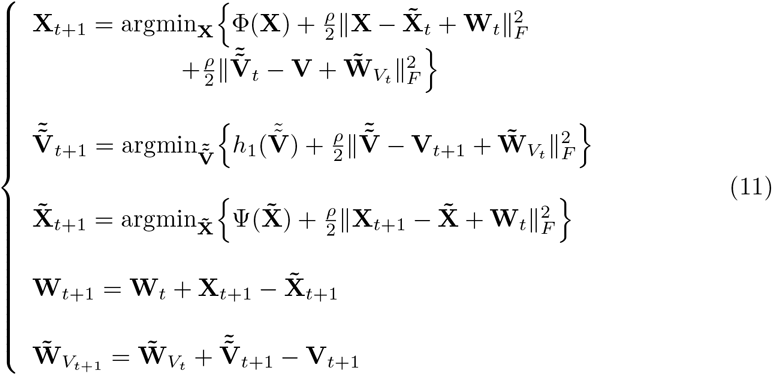

#### Theorem 1.

*There exists a solution* 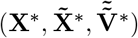 *to the EDOHA optimization problem (3), and the ADMM iterations via (11) approach the optimal value, i*.*e. p*_*t*_ → *p*^*∗*^, *where* 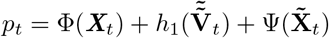 *and* 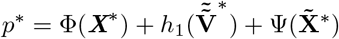.

The theorem establishes the convergence of the ADMM algorithm to achieve the optimal solution for EDOHA. It also automatically establishes algorithmic convergence for any optimization problem that can be regarded as a sub-case of EDOHA, for example, JRmGRN, which was not established before. A general algorithm for solving the optimization problem is shown in S1 Text. And the proof of Theorem 1 is shown in S2 Text.

### Faster computations for EDOHA

We now present a theorem that leads to substantial computational improvements to the EDOHA. Using the theorem, one can inspect the empirical covariance matrices **S**^(1)^, …, **S**^(*K*)^ in order to determine whether the solution to the EDOHA optimization problem is block diagonal after some permutation of the features. Previous studies [6, 7] use uniform thresholding to decompose the precision matrices of different classes in exactly the same way. Non-uniform thresholding generates a non-uniform feasible partition by thresholding the K empirical covariance matrices separately. In a non-uniform partition, two variables of the same group in one class may belong to different groups in another class [12]. Here we recommend a novel non-uniform thresholding approach that can split precision matrices into smaller submatrices without ignoring the different sparsity patterns from different matrices. Now we provide the key result. The following theorem states the sufficient conditions for the presence of non-uniform block diagonal structure.

#### Theorem 2.

*A sufficient condition for the solution to (2) to be block diagonal with blocks given by* 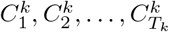 *is that*

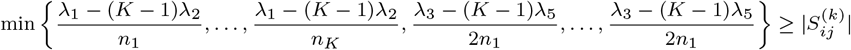

*for* 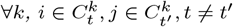.

Proof of Theorem 2 is given in S3 Text. Similar to Theorem 1 in [7], we decompose the reconstruction of a big network into the reconstruction of two or more small networks separately. JRmGRN has a sufficient condition for the presence of block diagonal structure. We now allow to split the precision matrices into class-specific block diagonal structures. It supplies us with a criterion if, given a partition of features 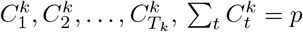, the solution of the optimization problem is block diagonal with each block corresponding to features in 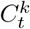. In practice, for any given (*λ*_1_, *λ*_2_, *λ*_3_, *λ*_4_, *λ*_5_), we can quickly perform the following two-step procedure to identify any block structure in each class in the solution.

- Create **B**^(*k*)^, a *p* ∗ *p* matrix with 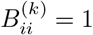 for *i* = 1, … , *p*. For *i* ≠ *j*, let 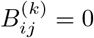 if the conditions specified in Theorem 2 are met for that pair of variables. Otherwise, set 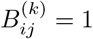.
- Identify the connected components of the undirected graph whose adjacency matrix is given by **B**^(*k*)^.

Theorem 2 guarantees that the connected components identified correspond to distinct blocks in *k*th class. Therefore, one can quickly obtain these solutions based on a non-uniform feasible partition. The block diagonal condition leads to massive computational speed-ups. Instead of computing th e eigen decomposition of *K p* ∗ *p* matrices, we compute the eigen decomposition of ∑_*k*_*T*_*k*_ matrices of dimensions 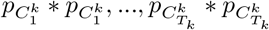. The computational complexity per-iteration is reduced from 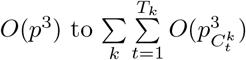.

### Tuning parameter selection

In this paper, we use Bayesian information criterion(BIC)-type quantity to select tuning parameters. We choose (*λ*_1_, *λ*_2_, *λ*_3_, *λ*_4_, *λ*_5_) to minimize the following function which balances the model likelihood and model complexity.

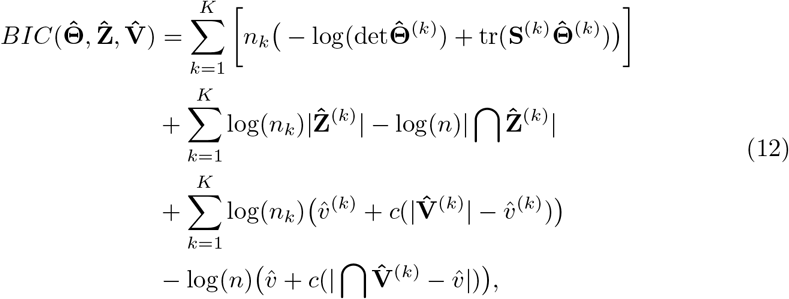

where 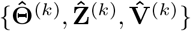 is the estimated parameters with a fixed set of tuning parameters (*λ*_1_, *λ*_2_, *λ*_3_, *λ*_4_, *λ*_5_), |∗| is the cardinality, 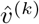 is the number of estimated hubs for *k*th class and 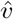 is the number of estimated common hubs, and *c* is a constant between zero and one. We select the set of tuning parameters (*λ*_1_, *λ*_2_, *λ*_3_, *λ*_4_, *λ*_5_) which minimizes the quality 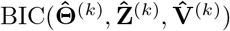. Note that BIC will favor more hub nodes in 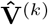 when constant *c* is small. In this paper, we take *c* = 0.3.

We use the grid search to find the tuning parameters. However, computing BIC over a range of values for five tuning parameters (*λ*_1_, *λ*_2_, *λ*_3_, *λ*_4_, *λ*_5_) may be computationally intensive. In this case, we suggest a dense search over (*λ*_1_, *λ*_3_, *λ*_4_) while holding (*λ*_2_, *λ*_5_) at fixed low values, followed by a quick search over (*λ*_2_, *λ*_5_), holding (*λ*_1_, *λ*_3_, *λ*_4_) at the selected values. With the number of features involved in the analysis dramatically increasing, tuning parameter selection becomes very complicated. In this situation, we need to explore some theoretical properties of the problem that can be used to provide guidance on our search of tuning parameters. This approach follows Deng et al. [7] and we provide the following theorems that extend their theoretical results to our present case with class-specific hubs.

#### Theorem 3.

*Let* (**Θ**^*∗*(*k*)^, ***Z***^*∗*(*k*)^, ***V***^*∗*(*k*)^)*be a solution to (2), a sufficient condition for* ***Z***^*∗*(*k*)^ *to be a diagonal matrix is that λ*_3_ + *λ*_4_ *<* 2*λ*_1_ *and λ*_5_ *<* 2*λ*_2_.

Proof of Theorem 3 is given in S4 Text.

#### Theorem 4.

*Let* (**Θ**^*∗*(*k*)^, ***Z***^*∗*(*k*)^, ***V***^*∗*(*k*)^)*be a solution to (2), a sufficient condition for* ***V***^*∗*(*k*)^ *to be a diagonal matrix is that* 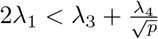 *and* 2*λ*_2_ *< λ*_5_.

Proof of Theorem 4 is given in S5 Text.

#### Corollary 1.

*Let* (**Θ**^*∗*(*k*)^, ***Z***^*∗*(*k*)^, ***V***^*∗*(*k*)^)*be a solution to (2), a necessary condition for both* ***Z***^*∗*(*k*)^ *and* ***V***^*∗*(*k*)^ *to be non-diagonal matrices is that tuning parameters satisfy any one of the following conditions:*

a. 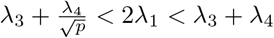
b. 2*λ*_2_ *< λ*_5_, 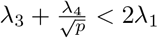
c. *λ*_5_ *<* 2*λ*_2_, 2*λ*_1_ *< λ*_3_ + *λ*_4_.

Specifically, we require that both **Z**^(*k*)^ and **V**^(*k*)^ are non-diagonal to produce non-trivial edges and hubs. With Corollary 1, we could reduce the search space of parameters *λ*_1_, *λ*_2_, *λ*_3_, *λ*_4_ and *λ*_5_ as these five tuning parameters are related. If *λ*_1_ and *λ*_2_ are large, and *λ*_3_, *λ*_4_ and *λ*_5_ are too small, then the elementary network **Z**^(*k*)^ may be very sparse and the number of hubs becomes huge. On the contrary, if *λ*_1_ and *λ*_2_ are quite small, and *λ*_3_, *λ*_4_ and *λ*_5_ are rather large, then we can get dense **Z**^(*k*)^ and few hubs. EDOHA’s conditions on tuning parameter selection are more complicated, since it involves *λ*_5_ which is not present for JRmGRN. In this paper, we use a uniformed grid of log space from 0.001 to 5 (size=20) for parameter *λ*_1_, *λ*_2_, *λ*_3_, *λ*_4_ and *λ*_5_ satisfying the conditions in Corollary 1.

### EDOHA for compositional data

Numerous studies have shown strong evidence that microbial compositions are closely related with various diseases such as diabetes [13], inflammatory bowel disease [14] and obesity [15]. Microbial count data are usually generated by sequencing variable regions of bacterial 16S rRNA gene. They are not directly comparable across samples and are usually transformed to relative abundance or proportion by dividing the total counts in the sample. A wide range of methods have been proposed to construct biological correlation networks for composition data, such as SPIEC-EASI [16], SparCC [17], Reboot [18], REBACCA [19], CCLasso [20] and COAT [21] for microbial interaction networks. However, these methods are for one class only.

To apply EDOHA to compositional data, we first perform data transformation. Here we briefly discuss compositional data for one class using microbiome data as an example. The absolute abundances or counts of *p* microbes, **y** = [*y*_1_, *y*_2_, … , *y*_*p*_], living in an environment such as human gut are usually not directly observable. However, the relative abundances 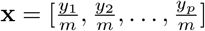 where 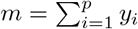, can be measured using 16S rRNA sequencing technologies. Here we apply the centered log-ratio transform [22] to remove the unit-sum constraint of compositional data. For a compositional variable **x** = (*x*_1_, … , *x*_*p*_), we have

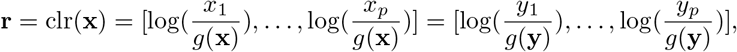

where 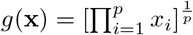 is the geometric mean of the composition vector. It is easy to show that there is a relationship between the covariance matrix **Σ** of **r** and the population covariance of the log-transformed absolute abundances 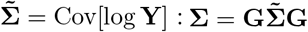 [16, 22], where 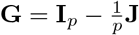, **I**_*p*_ is the *p*-dimensional identity matrix, and **J** is *p* ∗ *p* matrix with each of the entries equals 1. Kurtz et al. [16] mention that the matrix **G** is close to the identity matrix for high-dimensional data, and thus a finite sample estimator **S** of **Σ** may be as a good approximation of the empirical covariance of log **Y**. Actually, Cao et al. [21] have shown that **Σ** could be a proxy for 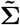 as long as 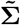 belongs to a class of large sparse covariance matrices. Therefore the interaction networks for high dimensional compositional data can be estimated based on the centered log-ratio transformed data.

## Results

### Simulation studies

To examine the efficiency of the proposed method for the better identification of common and class-specific hub nodes, we simulate Erd*ö*s-R*é*nyi (ER)-based network [23] and then generate corresponding compositional data to assess and validate the method. We compare the performance of EDOHA with the existing methods, such as the graphical lasso (JGL) and JRmGRN. Results show that EDOHA is more efficient than other methods when analyzing compositional data correlation networks which have both common and class-specific hub nodes.

#### Simulation strategy

To simulate a biological compositional data set such as microbiome count data, we consider the data are drawn with two steps. We first generate the basis abundance and proportion for each feature and then generate count data given a sequencing size (i.e. library size). The data structure characteristics are reflected in the basis covariance, which will be introduced in details later. Here we assume that basis proportions vary from sample to sample and are generated from one of three different distributions, namely, log ratio normal (LRN), Poisson log normal (LNP) and Dirichlet log normal (LND) distributions [19]. These three methods are presented in S6 Text. Then we extract count data from a multinomial distribution using the proportions, which reflects a random process that all sequences are equally likely to be selected in a biological sample.

To evaluate EDOHA comprehensively, we consider that the features are associated with ER-based network, in which each pair of nodes is selected with equal probability and connected with a predefined probability. The scale-free ER-based networks are generated by modifying the procedure used in Deng et al. [7]. Specifically, for a given number of classes (*K*), nodes (*p*), samples (*n*_*k*_), we use the following procedures to simulate ER-based network and corresponding compositional data.

**Step 1** We generate the base sparse matrix **A** in which *A*_*ij*_ is set as a random number in [−0.75, −0.25] ⋃ [0.25, 0.75] with probability *α* (elementary network sparsity 1-*α*) and zero otherwise.

**Step 2** Given the number of hubs *m*, we randomly choose *m* nodes and for each element that represents the correlation between *i*th hub node and other node *j*, 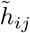, we set it to be a random number in [−0.75, −0.25] ⋃ [0.25, 0.75] with probability *β* (hub sparsity 1-*β*) and zero otherwise.

**Step 3** To construct the hub matrix **H**^(*k*)^, we randomly choose a fraction *δ* (network difference) of the hub nodes and reset them to be random numbers from 1, 2, … , *p*. The modified hub nodes are denoted by *h*^(*k*)^. As for nonzero elements in **H**^(*k*)^, we first set 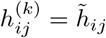 and then randomly adjust a fraction of *δ* of these nonzero elements and reset their values to be random numbers in [−0.75, −0.25] ⋃ [0.25, 0.75] with probability *β*.

**Step 4** To construct the elementary network, **Z**^(*k*)^, we first set it equal to **A**, and then randomly choose a fraction of *δ* of elements and reset their values to be random numbers in [−0.75, −0.25] ⋃ [0.25, 0.75] with probability *α* and zero otherwise. We set **Z**^(*k*)^ = **Z**^(*k*)^ + *t*(**Z**^(*k*)^) so that **Z**^(*k*)^ is symmetric.

**Step 5** We define the precision matrix **Θ**^(*k*)^ as **Z**^(*k*)^ + **H**^(*k*)^ + (**H**^(*k*)^)^*T*^. If **Θ**^(*k*)^ is not positive definite, we add the diagonal element of **Θ**^(*k*)^ by 0.1 −*λ*_min_(**Θ**^(*k*)^), where *λ*_min_(**Θ**^(*k*)^) is the minimum eigenvalue of **Θ**^(*k*)^.

**Step 6** We generate the compositional data of *n*_*k*_ samples for the *k*th class from a multinomial distribution using the proportion obtained from LRN with basis covariance (**Θ**^(*k*)^)^*−*1^.

Here the simulation studies are conducted for three classes with 40 or 80 samples for each class. The elementary network sparsity, the hub sparsity and the network difference are set as 0.98, 0.7, 0.2, respectively. We simulate three networks with 80, 160, 300 nodes, respectively. As we have mentioned, we use the BIC and the grid search to find the appropriate tuning parameters and model.

#### Simulation results

We consider simulated network described in the previous section with 80, 160, 300 nodes and estimate corresponding system with sample size n=40, n=80, respectively. The effects of EDOHA penalties vary with the sample size. To better present the simulation study results, we multiply the tuning parameters (*λ*_1_, *λ*_2_, *λ*_3_, *λ*_4_, *λ*_5_) by the sample size before performing the EDOHA.

We compare the performance of EDOHA and JRmGRN of identifying non-zero edges and class-specific edges. The results are computed averaging over 100 simulated data sets. We say that an edge (*i, j*) in the *k*th network is detected if the estimated association 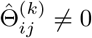 and we say that the edge is correctly detected if 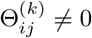. The number of differential edges, which differ between classes, is defined as follows [6]:

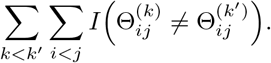

We record the sensitivity and specificity associated with detecting non-zero edges and detecting differential edges. The sensitivity is the proportion of the non-zero or differential edges that are correctly detected and the specificity represents the proportion of the zero or non-differential edges that are correctly detected. Hence the sensitivity and specificity of edge detection (ED) and differential edge detection (DED) are computed as

- 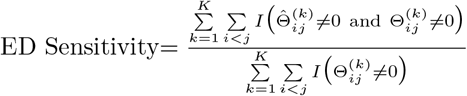
- 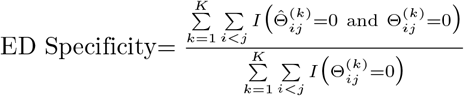
- 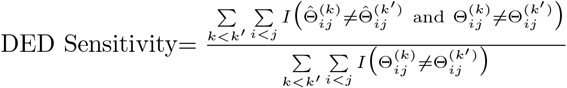
- 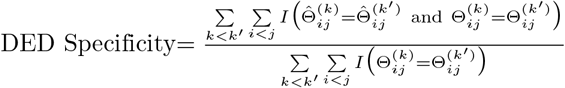

As shown in Table 1, if only the number of non-zero edges is considered, there is little difference between EDOHA and JRmGRN in terms of the total number of detected pairwise node-node associations. However, the sensitivity of detecting differential edges using EDOHA has more than doubled in all cases compared with JRmGRN. This is mainly because EDOHA is equipped with better ability to identify the class-specific edges.

**Table 1.** Means (Standard deviations) over 100 replicates using EDOHA and JRmGRN are shown for sensitivity and specificity of edge detection (ED) and differential edge detection (DED)

|  |  | n=40 |  |  |  | n=80 |  |  |  |
| --- | --- | --- | --- | --- | --- | --- | --- | --- | --- |
|  |  | ED | ED | DED | DED | ED | ED | DED | DED |
|  |  | Sensitivity | Specificity | Sensitivity | Specificity | Sensitivity | Specificity | Sensitivity | Specificity |
| p=80 | EDOHA | 0.614(0.089) | 0.925(0.054) | 0.396(0.098) | 0.959(0.049) | 0.597(0.110) | 0.960(0.024) | 0.416(0.104) | 0.968(0.019) |
|  | JRmGRN | 0.535(0.055) | 0.914(0.039) | 0.138(0.112) | 0.986(0.011) | 0.542(0.132) | 0.928(0.058) | 0.161(0.095) | 0.989(0.004) |
| p=160 | EDOHA | 0.352(0.055) | 0.974(0.011) | 0.288(0.059) | 0.977(0.005) | 0.430(0.063) | 0.977(0.010) | 0.329(0.058) | 0.979(0.006) |
|  | JRmGRN | 0.355(0.039) | 0.955(0.012) | 0.077(0.043) | 0.992(0.004) | 0.417(0.053) | 0.989(0.003) | 0.114(0.068) | 0.987(0.002) |
| p=300 | EDOHA | 0.347(0.032) | 0.971(0.007) | 0.217(0.081) | 0.974(0.016) | 0.288(0.036) | 0.990(0.005) | 0.293(0.050) | 0.978(0.011) |
|  | JRmGRN | 0.297(0.052) | 0.939(0.025) | 0.068(0.067) | 0.988(0.018) | 0.253(0.028) | 0.991(0.003) | 0.110(0.037) | 0.991(0.002) |

We then show that EDOHA has substantial improvements over several other methods. Since the hub nodes cannot be found out by JGL explictly, the precision recall curve is constructed based on the differential non-zero edges in the network, which is compared with the results from aforementioned methods intuitively. We simulate the networks with varying sparsity and similarity in two conditions and estimate corresponding networks with 160 nodes. The sample size is 80. To compare the results from different methods, we simulate each situation 100 times. As can be seen in Fig 1, the precision of EDOHA stays high through a larger range of recall, whereas for the other methods it quickly drops to the level of random guessing. This agrees with our expectation since EDOHA distinguishes the differences among elementary networks and hub networks respectively, which fits the data in the model better.

**Fig 1.**
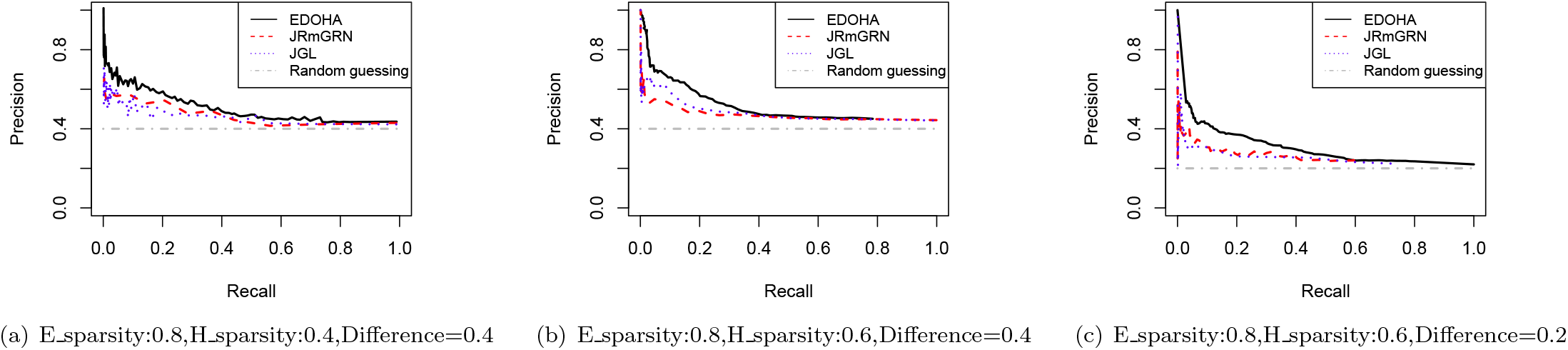
The Precision-Recall curve of EDOHA, JRmGRN and JGL for differential edge detection under different networks settings. ‘E _sparsity’ is the sparsity of elementary network; ‘H _sparsity’ is the sparsity of Hub network, the last parameter shown in title is the difference of two elementary networks.

Hubs are explicitly modeled by EDOHA and JRmGRN. We simulate the networks with both common and class-specific hubs and compare the results with JRmGRN. To better present the performance of identifying class-specific hubs, we also compare the hub detection capability with HGL [24], which only handle data from a single class. When applying HGL, networks are fitted for each class separately. The entire procedure is repeated 50 times. Comprehensive evaluation of EDOHA on identifying the common and class-specific hubs are presented in Table 2. The true positive rate (TPR), false positive rate (FPR) and Precision for common (C) hubs and class-specific (S) hubs are defined as

**Table 2.** Performances of EDOHA, JRmGRN and HGL for hub detection are compared by True Positive (TP), False Positive (FP) and Precision. The network difference are set as 0.3. The results are averaged over 50 simulations.

|  |  | TPR-C | FPR-C | Precision-C | TPR-S | FPR-S | Precision-S | TPR | FPR | Precision |
| --- | --- | --- | --- | --- | --- | --- | --- | --- | --- | --- |
| 80 nodes (5 hubs) | EDOHA | 0.896 | 0.021 | 0.886 | 0.751 | 0.056 | 0.728 | 0.841 | 0.027 | 0.824 |
|  | JRmGRN | 0.986 | 0.046 | 0.532 | NA | NA | NA | 0.810 | 0.049 | 0.688 |
|  | HGL | 0.784 | 0.016 | 0.781 | 0.695 | 0.168 | 0.407 | 0.835 | 0.051 | 0.619 |
| 160 nodes (8 hubs) | EDOHA | 0.793 | 0.004 | 0.837 | 0.837 | 0.035 | 0.776 | 0.861 | 0.005 | 0.897 |
|  | JRmGRN | 0.904 | 0.029 | 0.643 | NA | NA | NA | 0.727 | 0.029 | 0.674 |
|  | HGL | 0.621 | 0.006 | 0.738 | 0.767 | 0.038 | 0.566 | 0.737 | 0.023 | 0.663 |
| 300 nodes (12 hubs) | EDOHA | 0.789 | 0.001 | 0.879 | 0.878 | 0.018 | 0.756 | 0.895 | 0.001 | 0.863 |
|  | JRmGRN | 0.887 | 0.032 | 0.654 | NA | NA | NA | 0.705 | 0.011 | 0.706 |
|  | HGL | 0.537 | 0.001 | 0.819 | 0.713 | 0.023 | 0.553 | 0.793 | 0.021 | 0.589 |

NA: JRmGRN could not detect class-specific hubs.

- 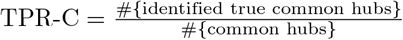
- 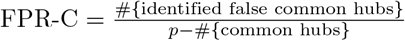
- 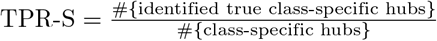
- 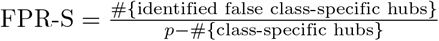
- 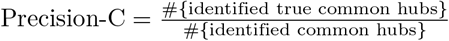
- 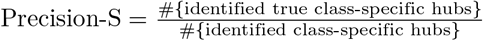

Total TPR, FPR and Precision are computed as

- 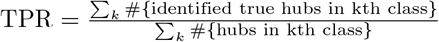
- 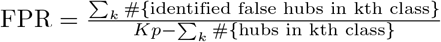
- 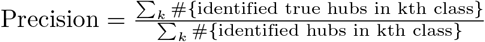

A simple example computing the TPR, FPR and Precision is described in S7 Text. Since JRmGRN only detects common hubs, there is no corresponding information of class-specific hubs. It can be seen that EDOHA has almost the highest precision and lowest FPR when we count the common hubs and class-specific hubs separately. Although JRmGRN works quite well in identifying common hubs, it tends to incorrectly identify some common hubs. As we mentioned earlier, JRmGRN can be viewed as a subcase of EDOHA, i.e. *λ*_5_ = ∞, and HGL is like EDOHA with *λ*_2_ = 0, *λ*_5_ = 0. Hence EDOHA has better performance than JRmGRN and HGL when analyzing correlation networks which have both common and class-specific hub nodes. Additional simulations for only common hubs and only class-specific hubs are shown in S1 Table. From simulation results, EDOHA could detect most common hubs in only common hubs setting and well recognize class-specific hubs in only class-specific hubs setting. We also find that the results of EDOHA and JRmGRN are similar to each other when most of true hubs are common ones but quite different when the true networks have more class-specific hubs. These results suggest the usefulness of EDOHA in identifying true hub nodes in a situation where one does not know if they are class-specific or common.

### Real data analysis

We apply the proposed model on three real data sets: one is proteomic data and the other two are microbiome data. Compared with the analysis methods used in the original publications, our model possesses the competence in constructing multiple networks with common and class-specific hubs across multiple classes. We also implement JRmGRN to infer interaction network and detect the hubs across classes. We find that some of hubs recognized by EDOHA, including common and class-specific ones, are identified as common hubs by JRmGRN. From simulation study, EDOHA may be more reliable when the results between them are significantly different.

#### Application to mouse skin microbiome data

We apply EDOHA to a mouse skin microbial data set (PRJEB1934) including three groups of individuals: non-immunized (Control), immunized-healthy (Healthy), and immunized-diseased (EBA). Microbial communities are measured utilizing variable regions of bacterial 16S rRNA sequencing data. These regions are amplified, sequenced, and then grouped into common Operational Taxonomic Units (OTUs) according to the similarity and quantified, with OTU counts serving as an intermediary to the underlying microbial populations abundances. The data set contains 131 core OTUs mainly coming from four prime phyla, which are Firmicutes (44 OTUs), Proteobacteria (35 OTUs), Bacteroidetes (26 OTUs), Actinobacteria (17 OTUs). We analyze their abundance data from 261 mouse skin samples. In particular, we wish to reconstruct the pair-wise conditional correlations network and identify the OTUs that are hubs. Such OTUs likely play an important role in the environment.

In Fig 2, we plot the networks for the three groups. The hub OTUs are highlighted in orange. Only OTUs from Firmicutes and Actinobacteria are identified as hub OTUs. The three networks share only one common hub while Healthy and EBA groups have another common hub. However, three hub OTUs in the Healthy group do not appear as hubs in the EBA group. Note that two OTU hubs shared by the Healthy and the Control groups are not hubs in the EBA group. Such information may be useful to understand the mechanism of protection from disease, which would not be available without our method of class-specific hub detection. In contrary, JRmGRN identifies eight common hubs, four of which are detected by EDOHA as class-specific hubs, one in EDOHA’s common hub. Only one hub recognized by EDOHA is not included in JRmGRN’s common hub set.

**Fig 2.**
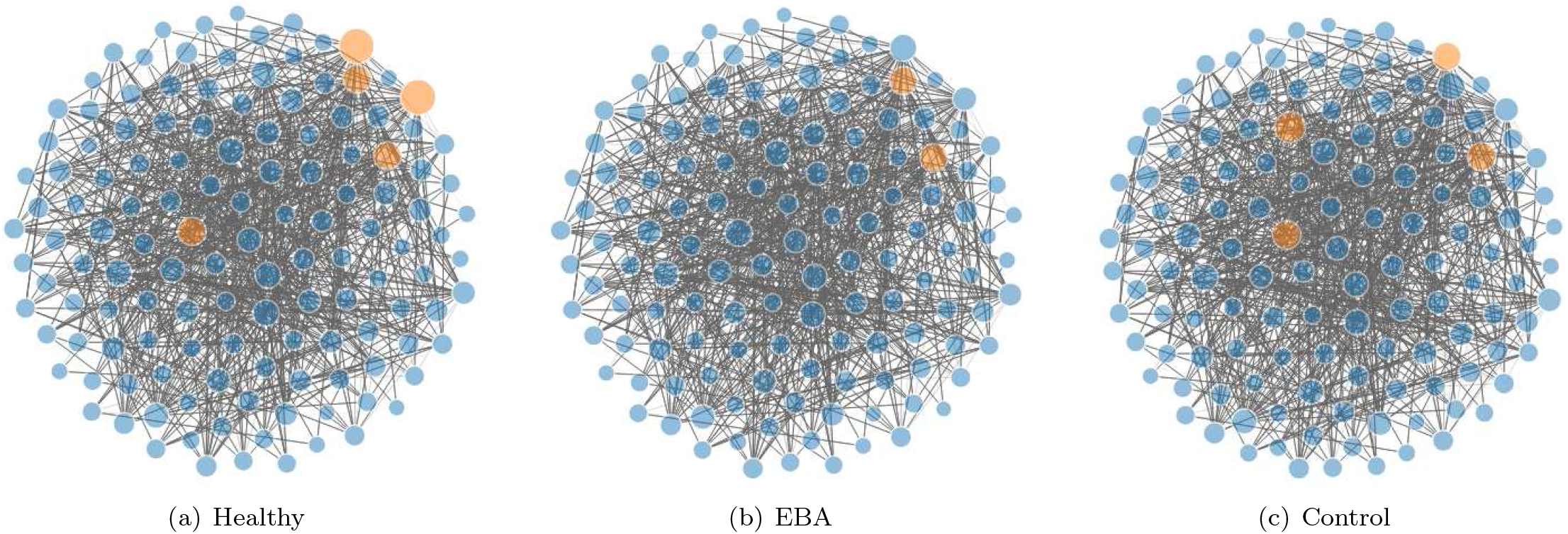
The estimated microbial networks for the groups of non-immunized (Control), immunized-healthy (Healthy) and immunized-diseased (EBA) individuals. The hub OTUs are highlighted in orange.

In addition to comparing the hub OTUs, we also investigate whether the correlation patterns among the 131 OTUs are different for different groups of disease status. A correlated pair of OTUs is considered consistent between two groups if the correlations in both groups have the same sign. We come to the same conclusion that correlations from the non-immunized individuals are less consistent with other two immunized groups than between the two immunized groups. There are 687 consistent pairs between the two immunized groups, while there are only 639 consistent pairs between the Control and Healthy and 632 between the Control and EBA groups. The results obtained in Ban et al. [19] were 532 consistent pairs between the two immunized groups, the other two were 236 and 212 respectively. Hence the gaps between these groups in our research are much less (See S1 Fig). The reason is mainly that we jointly model multiple networks simultaneously so that the similarity of network could be constructed more accurately by using datasets from multiple classes, which results in more accurate class-specific networks.

#### Application to IBD microbiome data

We preform our proposed method on the inflammatory bowel disease (IBD) multi-omics database from the Integrative Human Microbiome Project (HMP2 metadata) focusing on the functions of microbes in human health and disease. IBD further includes two main subtypes, Crohn’s disease (CD) and ulcerative colitis (UC). Our samples consist of 86 CD patients, 46 UC patients and 46 healthy controls with 342 OTUs. As is known, IBD is a chronic and relapsing inflammatory condition of the gastrointestinal (GI) tract and the GI microbiome of healthy humans is dominated by four major bacterial phyla: Firmicutes, Bacteroidetes, Proteobacteria and Actinobacteria. The data set contains 225, 44, 38, 23 OTUs from these four prime phyla respectively.

We aim to reconstruct the multiple microbial networks of the human gut that represent the interactions among the OTUs, as well as to identify hub OTUs that tend to have many interactions with other ones. Identifying such regulatory OTUs will lead to a better understanding of the mechanism of IBD, and eventually may lead to new therapeutic treatments. A large-scale cross-measurement type association network for host and microbial molecular interactions has been constructed [25]. Fig 3 displays the microbial interaction networks for the three classes. More hub OTUs are identified in CD than in UC and healthy controls. And almost each hub in UC and healthy groups is covered in the hub sets of CD group. We find that species from Actinobacteria are not detected as hub OTUs in three groups. Several studies [14, 26, 27] discovered that Faecalibacterium were differentially abundant in IBD and healthy group. Subdoligranulum, Roseburia and Fusobacterium have also been identified as hubs, all of which are associated, metatranscriptionally as well as metagenomically, with taxonomic features. In our study, Rumiococcus gnavus and Roseburia are found in CD and UC groups but not in healthy group, OTUs from Alistiles are only detected as hubs in CD group, which may lead to an entirely new line of medical research into IBD. By comparison, JRmGRN identifies thirteen common hubs, of which six are common ones, and six are shared in two classes, according to EDOHA. The remaining one is not found in EDOHA’s list of any class-specific hubs.

**Fig 3.**
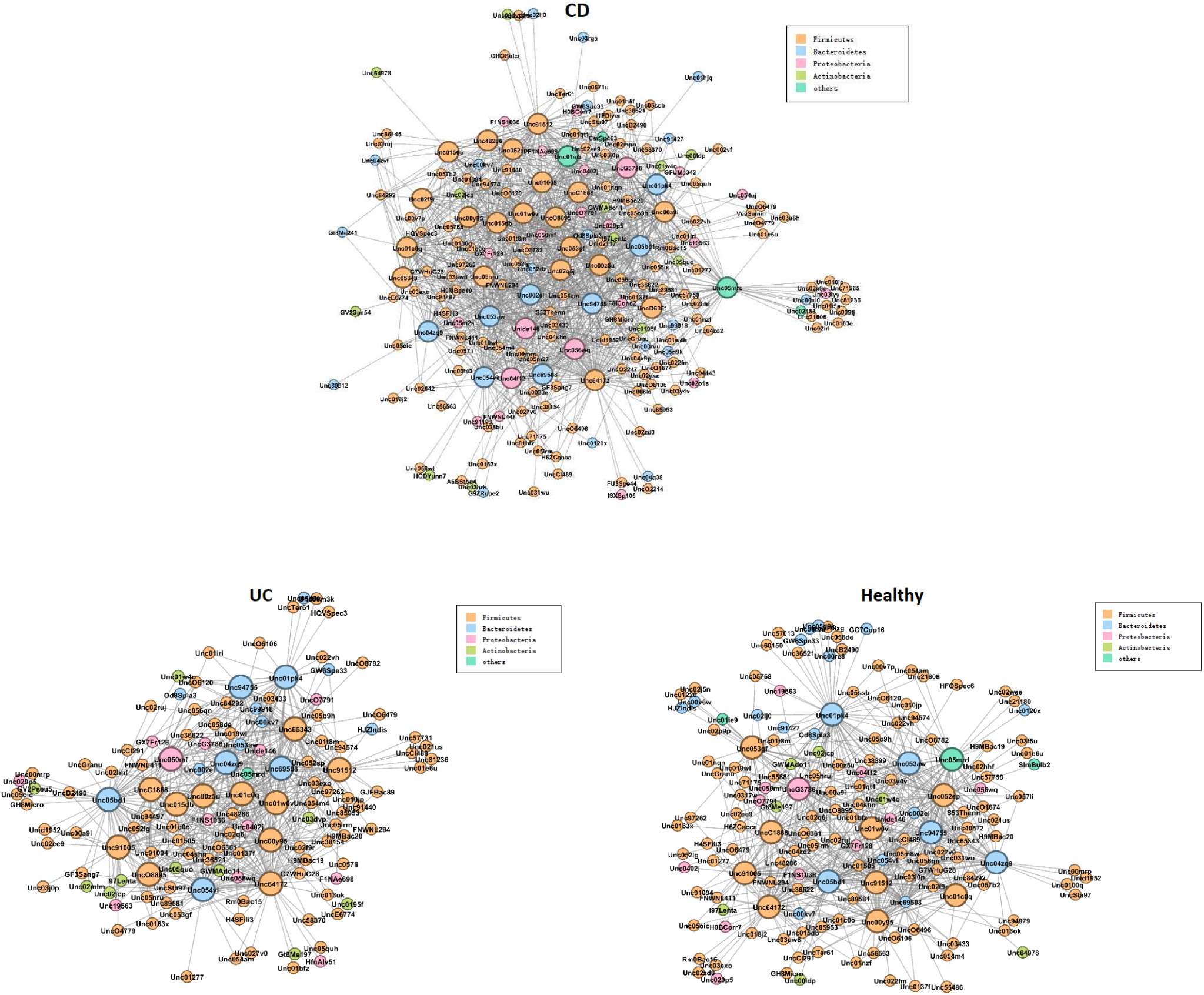
The estimated microbial networks for CD, UC and healthy groups. Four major bacterial phyla are marked by different colors. The larger nodes represent the hub OTUs.

#### Application to SARS-CoV-2 infection proteomic data

A most recent study have identified 332 high-confidence SARS-CoV-2 protein-human protein interactions that are connected with multiple biological processes [28]. In the 332 proteins interacted with SARS-CoV-2, 188 of them may interact with the major virus components. We search for the existence of the 188 proteins in four kinds of tissues: colon, liver, lung and kidney, and apply the proposed method, EDOHA, to construct proteome-wide networks and reveal common key hubs across different types of tissues and tissue-specific hubs. The proteomic data is downloaded from the National Cancer Institute Clinical Proteomic Tumor Analysis Consortium database (CPTAC).

As shown in Fig 4, we identify three common hub proteins DDX21, REEP6 and SEPSECS. And we identify MRPS5 as a hub only in colon, which is consistent with previous studies [8]. We also detect many other common hubs, including HMOX1, PRKAR2B and TIMM9, which appear as hubs in two or three organs. Moreover, BCKDK and COMT are involved in hubs only in liver. BWZ2, SLC44A2 and STOM are recognized as hubs merely in lung. And ATP1B1, ATP6AP1, ATP6V1A, CCDC86, ETFA, NUP210, PTGES2 and SCARB1 are screened as hubs only in kidney. All of these hub proteins detected in four tissues and their functions in living organism are shown in S2 Table. Eight hub proteins are recognized by JRmGRN. DDX21 and REEP6 are common hubs detected as common ones by EDOHA, while BZW2, CCDC86, MRPS5, PRKAR2B and STOM are class-specific hubs in one or two organs. RRP9 is the only one that is not in the list of EDOHA’s hub set.

**Fig 4.**
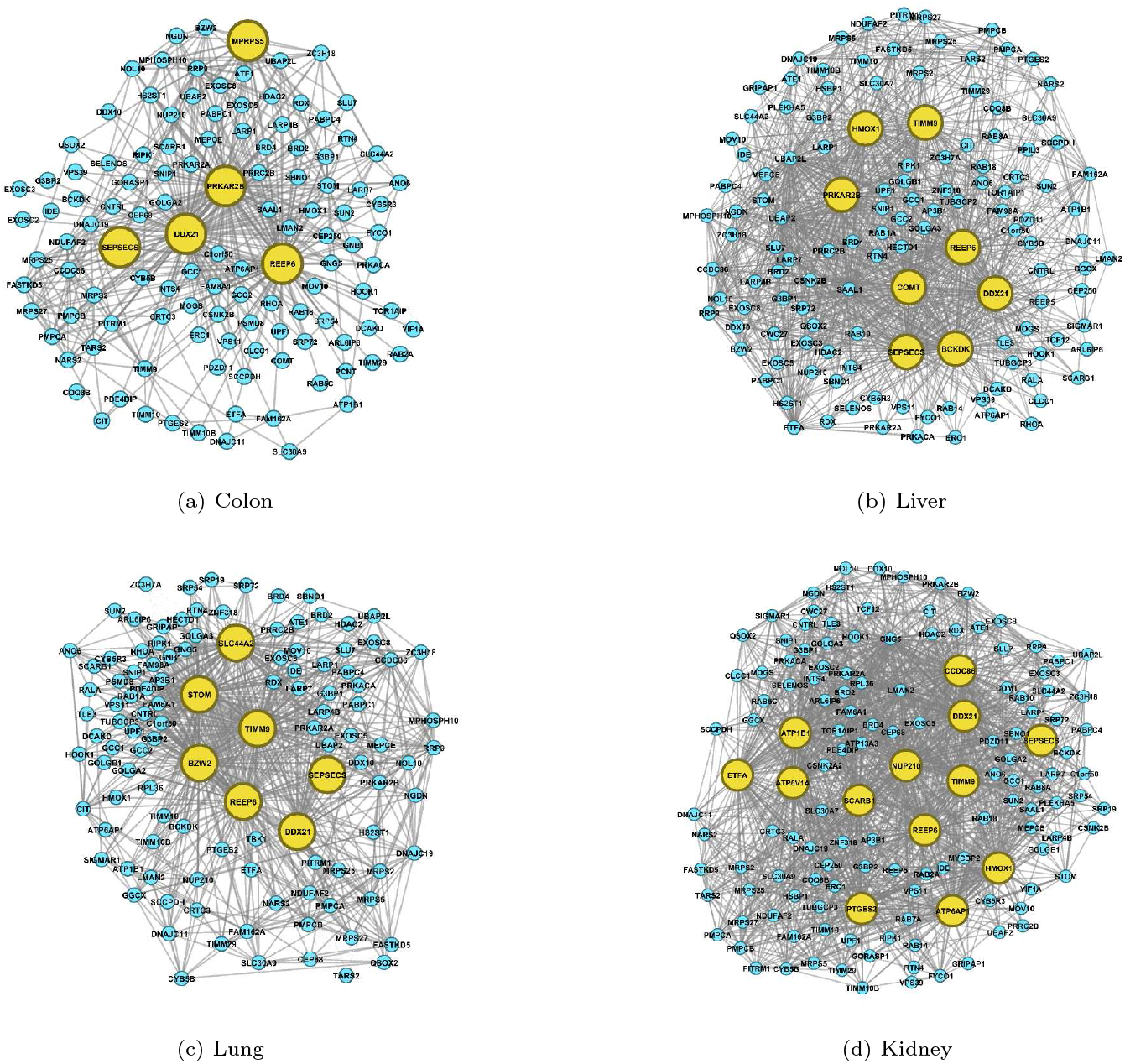
The estimated interaction networks of proteins affected by SARS-Cov-2 in four tissues. The hub proteins are highlighted in yellow.

All of these hubs have connectivity at least 4 times larger than that of any non-hubs. Ubiquitous hubs in multiple tissues would be promising drug targets to rescue multi-organ injury and deal with inflammation. Certain tissue-specific hubs might mediate specific dysfunction. Such information is urgently needed for the identification of the therapeutic targets for intervention and vaccine development.

## Discussion

Currently, there has been an increasing interest in the structure of multiple interaction networks. In most cases, people implicitly assume that each node has roughly the same number of interactions within the network when analyzing omics data, and each pair of nodes has equal probability to be an edge and all edges are independent of each other. However, this assumption is not appropriate in some real-world networks. In biological networks, scale-free properties are quite universal, which means the number of edges for each node follows a power-law distribution and a small proportion of nodes interact with many other ones. Barabasi and Oltvai [29] has found that most networks within the cell approximate a scale-free topology, including the metabolic networks, protein-protein interactions and genetic regulatory networks. The presence of hubs seems to be a general feature of all cellular networks. For example, hub proteins play critical roles in the organization and function of cellular protein interaction networks. It has also been demonstrated that such hub proteins may constitute an important pool of attractive drug targets. One typical aim is to capture more complex interactions and identify class-specific hubs in class-specific networks. Constructing biological association networks based on data sets from the same tissue with different phenotypes or different tissue enables us to screen out the influential features contributing to life health and disease, which provides insights into understanding the essential elements in living organisms and ecosystems. As researches into biological correlation networks continue, it has become important to develop a novel model to jointly estimate the scale-free interactions networks from different classes.

In this paper we propose a new statistical procedure to construct class-specific networks and select informative hub features among multiple classes for high dimensional omics data. Hub features, including common and specific ones, are accurately identified by decomposing the precision matrix into two parts. New penalty terms are added to single out class-specific hubs. Moreover, theoretical properties for selecting tuning parameters are investigated to improve computation efficiency. For a fixed set of tuning parameters, using a Mac desktop computer with 2.3 GHz Intel Core i5 processor and 8 GB 2133 MHz LPDDR3 memory, the average running times for estimating the precision matrices are about 2.5 min for 100 nodes, 7 min for 200 nodes, 20 min for 300 nodes, respectively. In future, we will explore strategies to speed up the computation, such as the randomized parameter search. The synthetic data are generated with ER-based network to model as closely as possible the situation in experimental biological compositional data. Our simulation studies show that the proposed method achieves higher accuracy in detecting the differential edges from different classes. We show that EDOHA has the potential to recognize the class-specific hub features and gains the larger area under the Precision-Recall curves compared with other methods. We also apply the proposed method on three real omics data sets. One of them is proteomic data from different tissues, and the other two are microbial data from microbial communities with different phenotypes. Across all three data sets, EDOHA successfully builds multiple networks and the results are basically consistent with previous reports. Furthermore, EDOHA identifies some hub features, both common and class-specific ones, which provides a deeper understanding of the mechanisms involved. Overall, EDOHA could not only jointly reconstruct multiple networks but also detect class-specific hubs explicitly for omics data with multiple distinct classes. It is promising in generating networks with such data structure.

EDOHA is in fact a general method applicable to many types of omics data such as gene expression data, which follow multivariate normal distribution. When EDOHA is applied to compositional data, one only needs to take the centered log-ratio transformed data as input. In fact, many other interaction network methods based on Gaussian graphical models have been proposed to account for compositionality more recently, such as gCoda [30], CD-trace [31], and BC-gLASSO [32]. One of our future work is to decompose the precision matrix in these newer methods as Θ = *Z* + *V* + (*V*)^*T*^ and use the penalty function *P* (Θ) in our method to construct multiple interaction networks with common and class-specific hubs.

## Supporting information

**S1 Text. A detailed ADMM algorithm for EDOHA**.

**S2 Text. The proof of the convergence of the ADMM algorithm for EDOHA**.

**S3 Text. The proof of the sufficient conditions for the non-uniform block diagonal structure**.

**S4 Text. The proof of Theorem 3**.

**S5 Text. The proof of Theorem 4**.

**S6 Text. Methods for generating the basis proportion for each feature**.

**S7 Text. A simple example computation of TPR and FPR based on current method**.

**S1 Table. Additional simulations for situations with only common hubs and with only class-specific hubs**.

**S1 Fig. Venn diagram of consistent correlated OTUs from Control, Healthy and EBA groups**. The figure shows the number of possible pairs within the same group and between different groups. It is suggested that the gaps between these groups in our research are much less than Ban et al. [19].

**S2 Table. The hub proteins detected in four organs and their functions in living organism**. The table lists hub proteins detected as common ones as well as tissue-specific ones, and introduces their functions.

## Acknowledgements

We thank the reviewers for their comments which are very helpful in improving our paper. We also thank Professor Olga Vitek for sharing her knowledge about the proteomic data.

