## supplemental files for "Extended Graphical Lasso for Multiple Interaction Networks for High Dimensional Omics Data"

### S1 Text: A detailed ADMM for the extended joint hub graphical lasso

Since the objective function is completely separable with respect to the variables $(\Theta^{(k)}, Z^{(k)}, V^{(k)})$, updating $X_{t+1}$ can be achieved by updating each variable.

$$\Theta_{t+1}^{(k)} = \text{argmin}\Big\{-n_k(\log(\det(\Theta^{(k)}))-\text{tr}(S^{(k)}\Theta^{(k)}))+\frac{\rho}{2}||\Theta^{(k)}-\tilde{\Theta}_t^{(k)}+W_{\Theta_t}^{(k)}||_F^2\Big\}. \tag{1}$$

Let $UDU^T$ denote the eigen decomposition of $S^{(k)} - \frac{\rho}{n_k}(\tilde{\Theta}_t^{(k)} - W_{\Theta_t}^{(k)})$, the solution is given by $U\hat{D}U^T$, where $\hat{D}$ is the diagonal matrix with $j^{th}$ diagonal element as follows:

$$\frac{n_k}{2\rho}(D_{jj} + \sqrt{D_{jj}^2 + \frac{4\rho}{n_k}})$$

$$\begin{aligned} Z_{t+1} = \text{argmin}\Big\{\lambda_1 \sum_{k=1}^{K} ||Z^{(k)} - diag(Z^{(k)})||_1 + \lambda_2 \sum_{k<k'} ||Z^{(k)} - Z^{(k')} \\ - diag(Z^{(k)} - Z^{(k')})||_1 + \sum_{k=1}^{K} \frac{\rho}{2}||Z^{(k)} - \tilde{Z}_t^{(k)} + W_{Z_t}^{(k)}||_F^2\Big\}. \end{aligned} \tag{2}$$

The objection function is completely separable with respect to each pair of matrix elements $\{(i,j), i \neq j\}$. Therefore, it is equivalent to solve separate optimization problems.

$$\begin{aligned} Z_{tij} = \text{argmin}\Big\{\lambda_1 \sum_{k=1}^{K} |Z_{ij}^{(k)}| + \lambda_2 \sum_{k<k'} |Z_{ij}^{(k)} - Z_{ij}^{(k')}| \\ + \sum_{k=1}^{K} \frac{\rho}{2}(Z_{ij}^{(k)} - \tilde{Z}_{tij}^{(k)} + W_{Z(tij)}^{(k)})^2\Big\}. \end{aligned} \tag{3}$$

This form belongs to a class of fused lasso problems[3].

$$\begin{aligned} V_{t+1}^{(k)} = \text{argmin}\Big\{\lambda_3||V^{(k)} - diag(V^{(k)})||_1 + \lambda_4||V^{(k)} - diag(V^{(k)})||_{1,2} \\ + \frac{\rho}{2}||V^{(k)} - \tilde{V}_t^{(k)} + W_{V_t}^{(k)}||_F^2 + \frac{\rho}{2}||\tilde{\tilde{V}}_t^{(k)} - V^{(k)} + \tilde{W}_{V_t}^{(k)}||_F^2\Big\}, \end{aligned} \tag{4}$$

the objective function is column separable. Let $(V_{t+1})_j$ be the $j$th column of matrix $V_{t+1}$, we solve

$$\begin{aligned}(V_{t+1})_j^{(k)} &= \text{argmin}\Big\{\lambda_3||V_j^{(k)}||_1 + \lambda_4||V_j^{(k)}||_2 + \frac{\rho}{2}||V_j^{(k)} - (\tilde{V}_t)_j^{(k)} + (W_{V_t})_j^{(k)}||_2^2 \\ &\quad + \frac{\rho}{2}||(\tilde{\tilde{V}}_t)_j^{(k)} - V_j^{(k)} + (\tilde{W}_{V_t})_j^{(k)}||_2^2\Big\} \\ &= \text{argmin}\Big\{\lambda_3||V_j^{(k)}||_1 + \lambda_4||V_j^{(k)}||_2 - \rho < V_j^{(k)}, (\tilde{V}_t)_j^{(k)} + (\tilde{\tilde{V}}_t)_j^{(k)} - (W_{V_t})_j^{(k)} + (\tilde{W}_{V_t})_j^{(k)} > \\ &\quad + \rho||V_j^{(k)}||_2^2 + \frac{\rho}{2}||(\tilde{V}_t)_j^{(k)} - (W_{V_t})_j^{(k)}||_2^2 + \frac{\rho}{2}||(\tilde{\tilde{V}}_t)_j^{(k)} + (\tilde{W}_{V_t})_j^{(k)}||_2^2\Big\}.\end{aligned} \tag{5}$$

It is equivalent to solve the optimization problem

$$(V_{t+1})_j^{(k)} = \text{argmin}\Big\{\lambda_3||V_j^{(k)}||_1 + \lambda_4||V_j^{(k)}||_2 + \rho||V_j^{(k)} - ((\tilde{V}_t)_j^{(k)} + (\tilde{\tilde{V}}_t)_j^{(k)} - (W_{V_t})_j^{(k)} + (\tilde{W}_{V_t})_j^{(k)})/2||_2^2\Big\}. \tag{6}$$

The result is given in [2].

$$\tilde{\tilde{V}}_{t+1} = \text{argmin}\Big\{\lambda_5 \sum_{k<k'} ||\tilde{\tilde{V}}^{(k)} - \tilde{\tilde{V}}^{(k')} - diag(\tilde{\tilde{V}}^{(k)} - \tilde{\tilde{V}}^{(k')})||_1 + \frac{\rho}{2}\sum_{k=1}^{K} ||\tilde{\tilde{V}}^{(k)} - V_{t+1}^{(k)} + \tilde{W}_{V_t}^{(k)}||_F^2\Big\}, \tag{7}$$

it is similar to the solution for Z.

Although the objective function is not separable with respect to the variables $(\tilde{\Theta}^{(k)}, \tilde{Z}^{(k)}, \tilde{V}^{(k)})$, updating $\tilde{X}_{t+1}$, can be achieved as follows:

$$\Pi_{t+1}^{(k)} = \frac{\rho}{6}\Big(\Theta_{t+1}^{(k)} + W_{\Theta_t}^{(k)} - Z_{t+1}^{(k)} - W_{Z_t}^{(k)} - V_{t+1}^{(k)} - W_{V_t}^{(k)} - t(V_{t+1}^{(k)} + W_{V_t}^{(k)})\Big) \tag{8}$$

$$\tilde{\Theta}^{(k)} = -\frac{\Pi_{t+1}^{(k)}}{\rho} + \Theta_{t+1}^{(k)} + W_{\Theta_t}^{(k)} \tag{9}$$

$$\tilde{Z}^{(k)} = \frac{\Pi_{t+1}^{(k)}}{\rho} + Z_{t+1}^{(k)} + W_{Z_t}^{(k)} \tag{10}$$

$$\tilde{V}^{(k)} = 2\frac{\Pi_{t+1}^{(k)}}{\rho} + V_{t+1}^{(k)} + W_{V_t}^{(k)} \tag{11}$$

***Derivation of (8)***

$$\begin{aligned}\tilde{X}_{t+1} &= \text{argmin}_{\tilde{X}}\Big\{\Psi(\tilde{X}) + \frac{\rho}{2}||X_{t+1} - \tilde{X} + W_t||_F^2\Big\}\\ &= \text{argmin}\Big\{\sum_{k=1}^{K} I\big(\tilde{\Theta}^{(k)} = \tilde{Z}^{(k)} + \tilde{V}^{(k)} + t(\tilde{V}^{(k)})\big) + \frac{\rho}{2}||X_{t+1} - \tilde{X} + W_t||_F^2\Big\}\\ &= \text{argmin}\Big\{\frac{\rho}{2}\sum_{k=1}^{K}||\tilde{\Theta}^{(k)} - (\Theta_{t+1}^{(k)} + W_{\Theta_t}^{(k)})||_F^2\\ &\quad + \frac{\rho}{2}\sum_{k=1}^{K}||\tilde{Z}^{(k)} - (Z_{t+1}^{(k)} + W_{Z_t}^{(k)})||_F^2 + \frac{\rho}{2}\sum_{k=1}^{K}||\tilde{V}^{(k)} - (V_{t+1}^{(k)} + W_{V_t}^{(k)})||_F^2\Big\},\end{aligned}$$

where $\tilde{\Theta}^{(k)} = \tilde{Z}^{(k)} + \tilde{V}^{(k)} + t(\tilde{V}^{(k)})$.

The Lagrange form of the above formula is

$$\begin{aligned}&L(\tilde{\Theta}^{(k)}, \tilde{Z}^{(k)}, \tilde{V}^{(k)}, \Pi^{(k)})\\ &= \frac{\rho}{2}\sum_{k=1}^{K}||\tilde{\Theta}^{(k)} - (\Theta_{t+1}^{(k)} + W_{\Theta_t}^{(k)})||_F^2 + \frac{\rho}{2}\sum_{k=1}^{K}||\tilde{Z}^{(k)} - (Z_{t+1}^{(k)} + W_{Z_t}^{(k)})||_F^2\\ &\quad + \frac{\rho}{2}\sum_{k=1}^{K}||\tilde{V}^{(k)} - (V_{t+1}^{(k)} + W_{V_t}^{(k)})||_F^2 + \sum_{k=1}^{K}\text{tr}\Big(\Pi^{(k)}\big(\tilde{\Theta}^{(k)} - (\tilde{Z}^{(k)} + \tilde{V}^{(k)} + t(\tilde{V}^{(k)}))\big)\Big)\end{aligned}$$

$$\begin{aligned}\frac{\partial L}{\partial\tilde{\Theta}^{(k)}} &= \rho\big(\tilde{\Theta}^{(k)} - (\Theta_{t+1}^{(k)} + W_{\Theta_t}^{(k)})\big) + \Pi^{(k)} = 0\\ \frac{\partial L}{\partial\tilde{Z}^{(k)}} &= \rho\big(\tilde{Z}^{(k)} - (Z_{t+1}^{(k)} + W_{Z_t}^{(k)})\big) - \Pi^{(k)} = 0\\ \frac{\partial L}{\partial\tilde{V}^{(k)}} &= \rho\big(\tilde{V}^{(k)} - (V_{t+1}^{(k)} + W_{V_t}^{(k)})\big) - 2\Pi^{(k)} = 0\\ \frac{\partial L}{\partial t(\tilde{V}^{(k)})} &= \rho\big(t(\tilde{V}^{(k)}) - t(V_{t+1}^{(k)} + W_{V_t}^{(k)})\big) - 2\Pi^{(k)} = 0\end{aligned}$$

For each k, the first equality subtracts the last three equalities and we get

$$\rho\big(Z_{t+1}^{(k)} + W_{Z_t}^{(k)} + V_{t+1}^{(k)} + W_{V_t}^{(k)} + t(V_{t+1}^{(k)} + W_{V_t}^{(k)}) - (\Theta_{t+1}^{(k)} + W_{\Theta_t}^{(k)})\big) + 6\Pi^{(k)} = 0$$

Therefore

$$\Pi_{t+1}^{(k)} = \frac{\rho}{6}\Big((\Theta_{t+1}^{(k)} + W_{\Theta_t}^{(k)}) - (Z_{t+1}^{(k)} + W_{Z_t}^{(k)}) - (V_{t+1}^{(k)} + W_{V_t}^{(k)}) - t(V_{t+1}^{(k)} + W_{V_t}^{(k)})\Big)$$

**S2 Text: The proof of the convergence of the ADMM algorithm for EDOHA**

The scaled augmented Lagrangian can be rewritten as

$$\begin{aligned} L_\rho(X, \tilde{X}, \tilde{\tilde{V}}, U, \tilde{U}_V) = \Phi(X) + h_1(\tilde{\tilde{V}}) + \Psi(\tilde{X}) + U^T(X - \tilde{X}) + \frac{\rho}{2}\|X - \tilde{X}\|_F^2 \\ + \tilde{U}_V^T(\tilde{\tilde{V}} - V) + \frac{\rho}{2}\|\tilde{\tilde{V}} - V\|_F^2 \end{aligned} \tag{12}$$

where $U = \rho W$ and $\tilde{U} = \rho \tilde{W}_V$. Let $(X^*, \tilde{X}^*, \tilde{\tilde{V}}^*)$ be a primal optimal and $(U^*, U_V^*)$ be a dual optimal point. According to convex optimization theory, KKT conditions guarantee that the optimal duality gap is zero [1], that is,

$$L_0(X^*, \tilde{X}^*, \tilde{\tilde{V}}^*, U, \tilde{U}_V) \le L_0(X^*, \tilde{X}^*, \tilde{\tilde{V}}^*, U^*, \tilde{U}_V^*) \le L_0(X, \tilde{X}, \tilde{\tilde{V}}, U^*, \tilde{U}_V^*)$$

holds for all $X, \tilde{X}, \tilde{\tilde{V}}, U, \tilde{U}_V$. Since $(X^*, \tilde{X}^*, \tilde{\tilde{V}}^*, U^*, \tilde{U}_V^*)$ is a saddle point for $L_0$, we have

$$L_0(X^*, \tilde{X}^*, \tilde{\tilde{V}}^*, U^*, \tilde{U}_V^*) \le L_0(X_{t+1}, \tilde{X}_{t+1}, \tilde{\tilde{V}}_{t+1}, U^*, \tilde{U}_V^*).$$

Using $X^* = \tilde{X}^*$ and $V^* = \tilde{\tilde{V}}^*$,

$$p^* - p_{t+1} \le U^{*T}(X_{t+1} - \tilde{X}_{t+1}) + \tilde{U}_V^{*T}(\tilde{\tilde{V}}_{t+1} - V_{t+1}). \tag{13}$$

By definition, $X_{t+1}$ minimizes $L_\rho(X, \tilde{X}_t, \tilde{\tilde{V}}_t, U_t, \tilde{U}_{V_t})$. The optimality condition is

$$0 \in \partial L_\rho(X_{t+1}, \tilde{X}_t, \tilde{\tilde{V}}_t, U_t, \tilde{U}_{V_t}).$$

Since it is separable with respect to $(\Theta_{t+1}, Z_{t+1}, V_{t+1})$, we have

$$0 \in \partial f(\Theta_{t+1}) + U_{\Theta_t} + \rho(\Theta_{t+1} - \tilde{\Theta}_t).$$

Since $U_{\Theta_{t+1}} = U_{\Theta_t} + \rho(\Theta_{t+1} - \tilde{\Theta}_{t+1})$, we can obtain

$$0 \in \partial f(\Theta_{t+1}) + U_{\Theta_{t+1}} + \rho(\tilde{\Theta}_{t+1} - \tilde{\Theta}_t).$$

This implies that $\Theta_{t+1}$ minimizes

$$f(\Theta) + (U_{\Theta_{t+1}} + \rho(\tilde{\Theta}_{t+1} - \tilde{\Theta}_t))^T\Theta.$$

It follows that

$$f(\Theta_{t+1}) + (U_{\Theta_{t+1}} + \rho(\tilde{\Theta}_{t+1} - \tilde{\Theta}_t))^T \Theta_{t+1} \leq f(\Theta^*) + (U_{\Theta_{t+1}} + \rho(\tilde{\Theta}_{t+1} - \tilde{\Theta}_t))^T \Theta^*$$

Similarly, we have

$$g(Z_{t+1}) + (U_{Z_{t+1}} + \rho(\tilde{Z}_{t+1} - \tilde{Z}_t))^T Z_{t+1} \leq g(Z^*) + (U_{Z_{t+1}} + \rho(\tilde{Z}_{t+1} - \tilde{Z}_t))^T Z^*,$$

$$\begin{aligned} & h(V_{t+1}) + (U_{V_{t+1}} - \tilde{U}_{V_{t+1}} + \rho(\tilde{V}_{t+1} - \tilde{V}_t + \tilde{\tilde{V}}_{t+1} - \tilde{\tilde{V}}_t))^T V_{t+1} \\ & \leq h(V^*) + (U_{V_{t+1}} - \tilde{U}_{V_{t+1}} + \rho(\tilde{V}_{t+1} - \tilde{V}_t + \tilde{\tilde{V}}_{t+1} - \tilde{\tilde{V}}_t))^T V^* \end{aligned}$$

$$h_2(\tilde{\tilde{V}}_{t+1}) + \tilde{U}_{V_{t+1}}^T \tilde{\tilde{V}}_{t+1} \leq h_2(\tilde{\tilde{V}}^*) + \tilde{U}_{V_{t+1}}^T \tilde{\tilde{V}}^*$$

$$\Psi(\tilde{X}_{t+1}) - U_{t+1}^T \tilde{X}_{t+1} \leq \Psi(\tilde{X}^*) - U_{t+1}^T \tilde{X}^*$$

Add these inequalities above, using $X^* = \tilde{X}^*$ and $V^* = \tilde{V}^*$, we obtain

$$\begin{aligned} p_{t+1} - p^* \leq\ & U_{t+1}^T(\tilde{X}_{t+1} - X_{t+1}) + \tilde{U}_{V_{t+1}}^T(V_{t+1} - \tilde{\tilde{V}}_{t+1}) \\ & \rho(\tilde{X}_{t+1} - \tilde{X}_t)^T(X^* - X_{t+1}) + \rho(\tilde{\tilde{V}}_{t+1} - \tilde{\tilde{V}}_t)^T(V^* - V_{t+1}) \end{aligned} \tag{14}$$

Adding (13) and (14),

$$\begin{aligned} & (U_{t+1} - U^*)^T(\tilde{X}_{t+1} - X_{t+1}) + \rho(\tilde{X}_{t+1} - \tilde{X}_t)^T(X^* - X_{t+1}) \\ & + (\tilde{U}_{V_{t+1}} - \tilde{U}_V^*)^T(V_{t+1} - \tilde{\tilde{V}}_{t+1}) + \rho(\tilde{\tilde{V}}_{t+1} - \tilde{\tilde{V}}_t)^T(V^* - V_{t+1}) \geq 0 \end{aligned} \tag{15}$$

We begin by the first two terms. Substituting $X^* - X_{t+1} = X^* - \tilde{X}_{t+1} + \tilde{X}_{t+1} - X_{t+1}$ gives

$$(U_{t+1} - U^*)^T(\tilde{X}_{t+1} - X_{t+1}) + \rho(\tilde{X}_{t+1} - \tilde{X}_t)^T(\tilde{X}_{t+1} - X_{t+1}) + \rho(\tilde{X}_{t+1} - \tilde{X}_t)^T(\tilde{X}^* - \tilde{X}_{t+1}) \tag{16}$$

For the first term of (16),

$$\begin{aligned} & (U_t + \rho(X_{t+1} - \tilde{X}_{t+1}) - U^*)^T(\tilde{X}_{t+1} - X_{t+1}) \\ & = (U_t - U^*)^T(\tilde{X}_{t+1} - X_{t+1}) - \frac{\rho}{2}\|\tilde{X}_{t+1} - X_{t+1}\|^2 - \frac{\rho}{2}\|\tilde{X}_{t+1} - X_{t+1}\|^2 \\ & = (U_t - U^*)^T(\tilde{X}_{t+1} - X_{t+1}) - \frac{\rho}{2}\|\frac{1}{\rho}(U_{t+1} - U_t)\|^2 - \frac{\rho}{2}\|\tilde{X}_{t+1} - X_{t+1}\|^2 \\ & = -\frac{1}{\rho}(U_t - U^*)^T(U_{t+1} - U^* - (U_t - U^*)) \\ & - \frac{1}{2\rho}\|U_{t+1} - U^* - (U_t - U^*)\|^2 - \frac{\rho}{2}\|\tilde{X}_{t+1} - X_{t+1}\|^2 \\ & = \frac{1}{2\rho}\|U_t - U^*\|^2 - \frac{1}{2\rho}\|U_{t+1} - U^*\|^2 - \frac{\rho}{2}\|\tilde{X}_{t+1} - X_{t+1}\|^2. \end{aligned}$$

For the last two terms of (16), taking $\frac{\rho}{2}\|\tilde{X}_{t+1} - X_{t+1}\|^2$ from above,

$$\begin{aligned}
&-\frac{\rho}{2}\|\tilde{X}_{t+1} - X_{t+1}\|^2 + \rho(\tilde{X}_{t+1} - \tilde{X}_t)^T(\tilde{X}_{t+1} - X_{t+1})\\
&+\rho(\tilde{X}_{t+1} - \tilde{X}_t)^T(\tilde{X}^* - \tilde{X}_t + \tilde{X}_t + \tilde{X}_{t+1})\\
&= -\frac{\rho}{2}\|\tilde{X}_{t+1} - X_{t+1} - (\tilde{X}_{t+1} - \tilde{X}_t)\|^2 + \rho(\tilde{X}_{t+1} - \tilde{X}_t)(\tilde{X}^* - \tilde{X}_t) - \frac{\rho}{2}\|\tilde{X}_{t+1} - \tilde{X}_t\|^2\\
&= -\frac{\rho}{2}\|\tilde{X}_{t+1} - X_{t+1} - (\tilde{X}_{t+1} - \tilde{X}_t)\|^2 - \frac{\rho}{2}\|\tilde{X}_{t+1} - \tilde{X}^*\|^2 + \frac{\rho}{2}\|\tilde{X}_t - \tilde{X}^*\|^2
\end{aligned}$$

Hence the first two terms of (15) can be rewritten as

$$\frac{1}{2\rho}\|U_t - U^*\|^2 - \frac{1}{2\rho}\|U_{t+1} - U^*\|^2 - \frac{\rho}{2}\|\tilde{X}_{t+1} - X_{t+1} - (\tilde{X}_{t+1} - \tilde{X}_t)\|^2 + \frac{\rho}{2}\|\tilde{X}_t - \tilde{X}^*\|^2 - \frac{\rho}{2}\|\tilde{X}_{t+1} - \tilde{X}^*\|^2$$

Similarly, the last two terms of (15) can be rewritten as

$$\frac{1}{2\rho}\|\tilde{U}_{V_t} - \tilde{U}_V^*\|^2 - \frac{1}{2\rho}\|\tilde{U}_{V_{t+1}} - \tilde{U}_V^*\|^2 - \frac{\rho}{2}\|\tilde{\tilde{V}}_{t+1} - V_{t+1} - (\tilde{\tilde{V}}_{t+1} - \tilde{\tilde{V}}_t)\|^2 + \frac{\rho}{2}\|V_t - \tilde{\tilde{V}}^*\|^2 - \frac{\rho}{2}\|\tilde{\tilde{V}}_{t+1} - \tilde{\tilde{V}}^*\|^2$$

This implies that (15) can be written as

$$\begin{aligned}
&\frac{1}{2\rho}\|U_t - U^*\|^2 + \frac{1}{2\rho}\|\tilde{U}_{V_t} - \tilde{U}_V^*\|^2 + \frac{\rho}{2}\|\tilde{X}_t - \tilde{X}^*\|^2 + \frac{\rho}{2}\|V_t - \tilde{\tilde{V}}^*\|^2\\
&-\frac{1}{2\rho}\|U_{t+1} - U^*\|^2 - \frac{1}{2\rho}\|\tilde{U}_{V_{t+1}} - \tilde{U}_V^*\|^2 - \frac{\rho}{2}\|\tilde{X}_{t+1} - \tilde{X}^*\|^2 - \frac{\rho}{2}\|\tilde{\tilde{V}}_{t+1} - \tilde{\tilde{V}}^*\|^2\\
&-\frac{\rho}{2}\|\tilde{X}_{t+1} - X_{t+1} - (\tilde{X}_{t+1} - \tilde{X}_t)\|^2 - \frac{\rho}{2}\|\tilde{\tilde{V}}_{t+1} - V_{t+1} - (\tilde{\tilde{V}}_{t+1} - \tilde{\tilde{V}}_t)\|^2 \geq 0.
\end{aligned} \tag{17}$$

Let

$$D_t = \frac{1}{2\rho}\|U_t - U^*\|^2 + \frac{1}{2\rho}\|\tilde{U}_{V_t} - \tilde{U}_V^*\|^2 + \frac{\rho}{2}\|\tilde{X}_t - \tilde{X}^*\|^2 + \frac{\rho}{2}\|V_t - \tilde{\tilde{V}}^*\|^2$$

it follows that

$$D_t - D_{t+1} \geq \frac{\rho}{2}\|\tilde{X}_{t+1} - X_{t+1} - (\tilde{X}_{t+1} - \tilde{X}_t)\|^2 + \frac{\rho}{2}\|\tilde{\tilde{V}}_{t+1} - V_{t+1} - (\tilde{\tilde{V}}_{t+1} - \tilde{\tilde{V}}_t)\|^2 \tag{18}$$

Recall that $\tilde{X}_{t+1}$ minimizes $\Psi(\tilde{X}) - U_{t+1}^T\tilde{X}$ and $\tilde{X}_t$ minimizes $\Psi(\tilde{X}) - U_t^T\tilde{X}$, so we can add

$$\Psi(\tilde{X}_{t+1}) - U_{t+1}^T\tilde{X}_{t+1} \leq \Psi(\tilde{X}_t) - U_{t+1}^T\tilde{X}_t$$

and

$$\Psi(\tilde{X}_t) - U_t^T\tilde{X}_t \leq \Psi(\tilde{X}_{t+1}) - U_t^T\tilde{X}_{t+1}$$

to get

$$(U_{t+1} - U_t)^T (\tilde{X}_{t+1} - \tilde{X}_t) \geq 0.$$

Hence by $\rho > 0$,

$$(\tilde{X}_{t+1} - X_{t+1})^T (\tilde{X}_{t+1} - \tilde{X}_t) \leq 0.$$

Likewise,

$$(\tilde{\tilde{V}}_{t+1} - V_{t+1})^T (\tilde{\tilde{V}}_{t+1} - \tilde{\tilde{V}}_t) \leq 0.$$

Combined with (18),

$$D_t - D_{t+1} \geq \frac{\rho}{2}\|\tilde{X}_{t+1} - X_{t+1}\|^2 + \frac{\rho}{2}\|\tilde{X}_{t+1} - \tilde{X}_t\|^2 + \frac{\rho}{2}\|\tilde{\tilde{V}}_{t+1} - V_{t+1}\|^2 + \frac{\rho}{2}\|\tilde{\tilde{V}}_{t+1} - \tilde{\tilde{V}}_t\|^2.$$

Iterating these inequalities gives that

$$\sum_{t=0}^{\infty} \frac{\rho}{2}\|\tilde{X}_{t+1} - X_{t+1}\|^2 + \frac{\rho}{2}\|\tilde{X}_{t+1} - \tilde{X}_t\|^2 + \frac{\rho}{2}\|\tilde{\tilde{V}}_{t+1} - V_{t+1}\|^2 + \frac{\rho}{2}\|\tilde{\tilde{V}}_{t+1} - \tilde{\tilde{V}}_t\|^2 \leq D_0$$

which implies that $\tilde{X}_{t+1} - X_{t+1} \to 0$ and $\tilde{\tilde{V}}_{t+1} - V_{t+1} \to 0$. The right hand in (13) goes to zero as $t \to \infty$. Hence we have $\lim_{t\to\infty} p_t = p^*$.

### S3 Text: The proof of the sufficient conditions for the non-uniform block diagonal structure

Let $\Omega_k = \bigcup_{t=1}^{T_k}\{C_t^k * C_t^k\}, S_{\max}^{(k)} = \max_{(i,j)\in\Omega_k^c}|S_{ij}^{(k)}|$, let $A_{\Omega_k}$ denote the restriction of the matrix A to the set $\Omega_k$, that is

$$(A_{\Omega_k})_{ij} = \begin{cases} A_{ij}, & \text{if } (i,j) \in \Omega_k; \\ 0, & \text{if } (i,j) \notin \Omega_k. \end{cases}$$

Assume $(\{\Theta^{(k)}\}, \{Z^{(k)}\}, \{V^{(k)}\})$ is a feasible solution, then $(\{\Theta_{\Omega_k}^{(k)}\}, \{Z_{\Omega_k}^{(k)}\}, \{V_{\Omega_k}^{(k)}\})$ is also a feasible solution. We want to show that if the condition holds, then the objective value evaluated at $(\{\Theta_{\Omega_k}^{(k)}\}, \{Z_{\Omega_k}^{(k)}\}, \{V_{\Omega_k}^{(k)}\})$ is smaller than the objective value evaluated at $(\{\Theta^{(k)}\}, \{Z^{(k)}\}, \{V^{(k)}\})$. By Fischer's inequality,

$$-\text{logdet}(\Theta^{(k)}) > -\text{logdet}(\Theta_{\Omega_k}^{(k)})$$

We need only to prove

$$\sum_{k=1}^{K} n_k \mathrm{tr}(S^{(k)}\Theta^{(k)}) + \lambda_1 \sum_{k=1}^{K} ||Z^{(k)} - \mathrm{diag}(Z^{(k)})||_1 + \lambda_3 \sum_{k=1}^{K} ||V^{(k)} - \mathrm{diag}(V^{(k)})||_1$$
$$+ \lambda_2 \sum_{k<k'} ||Z^{(k)} - Z^{(k')} - \mathrm{diag}(Z^{(k)} - Z^{(k')})||_1 + \lambda_4 \sum_{k=1}^{K} ||V^{(k)} - \mathrm{diag}(V^{(k)})||_{1,2}$$
$$+ \lambda_5 \sum_{k<k'} ||V^{(k)} - V^{(k')} - \mathrm{diag}(V^{(k)} - V^{(k')})||_1$$
$$\geq \sum_{k=1}^{K} n_k \mathrm{tr}(S^{(k)}\Theta^{(k)}_{\Omega_k}) + \lambda_1 \sum_{k=1}^{K} ||Z^{(k)}_{\Omega_k} - \mathrm{diag}(Z^{(k)}_{\Omega_k})||_1 + \lambda_3 \sum_{k=1}^{K} ||V^{(k)}_{\Omega_k} - \mathrm{diag}(V^{(k)}_{\Omega_k})||_1$$
$$+ \lambda_2 \sum_{k<k'} ||Z^{(k)}_{\Omega_k} - Z^{(k')}_{\Omega_{k'}} - \mathrm{diag}(Z^{(k)}_{\Omega_k} - Z^{(k')}_{\Omega_{k'}})||_1 + \lambda_4 \sum_{k=1}^{K} ||V^{(k)}_{\Omega_k} - \mathrm{diag}(V^{(k)}_{\Omega_k})||_{1,2}$$
$$+ \lambda_5 \sum_{k<k'} ||V^{(k)}_{\Omega_k} - V^{(k')}_{\Omega_{k'}} - \mathrm{diag}(V^{(k)}_{\Omega_k} - V^{(k')}_{\Omega_{k'}})||_1 \tag{19}$$

As

$$\sum_{k<k'} ||Z^{(k)} - Z^{(k')}||_1 - \sum_{k<k'} ||Z^{(k)}_{\Omega_k} - Z^{(k')}_{\Omega_{k'}}||_1$$
$$= ||Z^{(k)}_{\Omega^c_k \bigcup \Omega^c_{k'}} - Z^{(k')}_{\Omega^c_k \bigcup \Omega^c_{k'}}||_1 - ||Z^{(k)}_{\Omega_k \bigcap \Omega^c_{k'}}||_1 - ||Z^{(k')}_{\Omega^c_k \bigcap \Omega_{k'}}||_1,$$
$$||V^{(k)}||_{1,2} \geq ||V^{(k)}_{\Omega_k}||_{1,2},$$
$$\sum_{k<k'} ||V^{(k)} - V^{(k')}||_1 - \sum_{k<k'} ||V^{(k)}_{\Omega_k} - V^{(k')}_{\Omega_{k'}}||_1$$
$$= ||V^{(k)}_{\Omega^c_k \bigcup \Omega^c_{k'}} - V^{(k')}_{\Omega^c_k \bigcup \Omega^c_{k'}}||_1 - ||V^{(k)}_{\Omega_k \bigcap \Omega^c_{k'}}||_1 - ||V^{(k')}_{\Omega^c_k \bigcap \Omega_{k'}}||_1,$$

Hence we only need to prove

$$\sum_{k=1}^{K} n_k < S^{(k)}_{\Omega^c_k}, \Theta^{(k)}_{\Omega^c_k} > + \lambda_1 \sum_{k=1}^{K} ||Z^{(k)}_{\Omega^c_k}||_1$$
$$+ \lambda_2 \sum_{k<k'} (||Z^{(k)}_{\Omega^c_k \bigcup \Omega^c_{k'}} - Z^{(k')}_{\Omega^c_k \bigcup \Omega^c_{k'}}||_1 - ||Z^{(k)}_{\Omega_k \bigcap \Omega^c_{k'}}||_1 - ||Z^{(k')}_{\Omega^c_k \bigcap \Omega_{k'}}||_1)$$
$$+ \lambda_3 \sum_{k=1}^{K} ||V^{(k)}_{\Omega^c_k}||_1 + \lambda_5 \sum_{k<k'} (||V^{(k)}_{\Omega^c_k \bigcup \Omega^c_{k'}} - V^{(k')}_{\Omega^c_k \bigcup \Omega^c_{k'}}||_1 - ||V^{(k)}_{\Omega_k \bigcap \Omega^c_{k'}}||_1 - ||V^{(k')}_{\Omega^c_k \bigcap \Omega_{k'}}||_1) \geq 0$$

As $||Z^{(k)}_{\Omega_k \bigcap \Omega^c_{k'}}||_1 \leq ||Z^{(k)}_{\Omega_k}||_1$ and $||V^{(k)}_{\Omega_k \bigcap \Omega^c_{k'}}||_1 \leq ||V^{(k)}_{\Omega_k}||_1$, we need to prove that

$$\sum_{k=1}^{K} n_k < S^{(k)}_{\Omega^c_k}, \Theta^{(k)}_{\Omega^c_k} > + (\lambda_1 - (K-1)\lambda_2) \sum_{k=1}^{K} ||Z^{(k)}_{\Omega^c_k}||_1 + (\lambda_3 - (K-1)\lambda_5) \sum_{k=1}^{K} ||V^{(k)}_{\Omega^c_k}||_1 \geq 0$$

$$
\begin{aligned}
&\sum_{k=1}^{K} n_k | < S^{(k)}_{\Omega_k^c}, \Theta^{(k)}_{\Omega_k^c} > | \\
= &\sum_{k=1}^{K} n_k | < S^{(k)}_{\Omega_k^c}, Z^{(k)}_{\Omega_k^c} + V^{(k)}_{\Omega_k^c} + (V^{(k)}_{\Omega_k^c})^T > | \\
\leq &\sum_{k=1}^{K} n_k | < S^{(k)}_{\Omega_k^c}, Z^{(k)}_{\Omega_k^c} > | + 2 \sum_{k=1}^{K} n_k | < S^{(k)}_{\Omega_k^c}, V^{(k)}_{\Omega_k^c} > | \\
\leq &\sum_{k=1}^{K} n_k S^{(k)}_{\max} ||Z^{(k)}_{\Omega_k^c}||_1 + 2 \sum_{k=1}^{K} n_k S^{(k)}_{\max} ||V^{(k)}_{\Omega_k^c}||_1 \\
\leq &(\lambda_1 + \lambda_2 - K\lambda_2) \sum_{k=1}^{K} ||Z^{(k)}_{\Omega_k^c}||_1 + (\lambda_3 + \lambda_5 - K\lambda_5) \sum_{k=1}^{K} ||V^{(k)}_{\Omega_k^c}||_1
\end{aligned}
$$

where the last inequality follows from the sufficient condition.

#### S4 Text: The proof of Theorem 3

Let $(\Theta^{*(k)}, Z^{*(k)}, V^{*(k)})$ be the solution to problem and suppose that $Z^{*(k)}$ is not a diagonal matrix. Let $\hat{Z}^{(k)} = \text{diag}(Z^{*(k)})$, and construct $\hat{V}$ as follows

$$
\hat{V}^{(k)}_{ij} = \begin{cases} V^{*(k)}_{ij} + \frac{Z^{*(k)}_{ij}}{2} & \text{if} \quad i \neq j \\ V^{*(k)}_{ij} & \text{otherwise} \end{cases}
$$

Then we have $\Theta^{*(k)} = \hat{Z}^{(k)} + \hat{V}^{(k)} + t(\hat{V}^{(k)})$. Thus $(\Theta^{*(k)}, \hat{Z}^{(k)}, \hat{V}^{(k)})$ is also a feasible solution. Now we prove that if the condition holds, $(\Theta^{*(k)}, \hat{Z}^{(k)}, \hat{V}^{(k)})$

has a smaller objective than $(\Theta^{*(k)}, Z^{*(k)}, V^{*(k)})$. We need to prove that

$$
\begin{aligned}
&\lambda_3 \sum_{k=1}^{K} ||\hat{V}^{(k)} - \mathrm{diag}(\hat{V}^{(k)})||_1 + \lambda_4 \sum_{k=1}^{K} ||\hat{V}^{(k)} - \mathrm{diag}(\hat{V}^{(k)})||_{1,2} \\
&+ \lambda_5 \sum_{k<k'} ||\hat{V}^{(k)} - \hat{V}^{(k')} - \mathrm{diag}(\hat{V}^{(k)} - \hat{V}^{(k')})||_1 \\
\le\ &\lambda_1 \sum_{k=1}^{K} ||Z^{*(k)} - \mathrm{diag}(Z^{*(k)})||_1 \\
&+ \lambda_2 \sum_{k<k'} ||Z^{*(k)} - Z^{*(k')} - \mathrm{diag}(Z^{*(k)} - Z^{*(k')})||_1 \\
&+ \lambda_3 \sum_{k=1}^{K} ||V^{*(k)} - \mathrm{diag}(V^{*(k)})||_1 + \lambda_4 \sum_{k=1}^{K} ||V^{*(k)} - \mathrm{diag}(V^{*(k)})||_{1,2} \\
&+ \lambda_5 \sum_{k<k'} ||V^{*(k)} - V^{*(k')} - \mathrm{diag}(V^{*(k)} - V^{*(k')})||_1.
\end{aligned}
\tag{20}
$$

Three terms in the left hand of formula (20) can be restricted separately.

$$
\begin{aligned}
&\lambda_3 \sum_{k=1}^{K} ||\hat{V}^{(k)} - \mathrm{diag}(\hat{V}^{(k)})||_1 \\
&\le \lambda_3 \sum_{k=1}^{K} ||V^{*(k)} - \mathrm{diag}(V^{*(k)})||_1 + \frac{\lambda_3}{2} \sum_{k=1}^{K} ||Z^{*(k)} - \mathrm{diag}(Z^{*(k)})||_1.
\end{aligned}
$$

$$
\begin{aligned}
&\lambda_4 \sum_{k=1}^{K} ||\hat{V}^{(k)} - \mathrm{diag}(\hat{V}^{(k)})||_{1,2} \\
&\le \lambda_4 \sum_{k=1}^{K} ||V^{*(k)} - \mathrm{diag}(V^{*(k)})||_{1,2} + \frac{\lambda_4}{2} \sum_{k=1}^{K} ||Z^{*(k)} - \mathrm{diag}(Z^{*(k)})||_{1,2}.
\end{aligned}
$$

$$
\begin{aligned}
&\lambda_5 \sum_{k<k'} ||\hat{V}^{(k)} - \hat{V}^{(k')} - \mathrm{diag}(\hat{V}^{(k)} - \hat{V}^{(k')})||_1 \\
&\le \lambda_5 \sum_{k<k'} ||V^{*(k)} - V^{*(k')} - \mathrm{diag}(V^{*(k)} - V^{*(k')})||_1 \\
&+ \frac{\lambda_5}{2} \sum_{k<k'} ||Z^{*(k)} - Z^{*(k')} - \mathrm{diag}(Z^{*(k)} - Z^{*(k')})||_1.
\end{aligned}
$$

Then the inequality (20) follows from the sufficient condition.

**S5 Text: The proof of Theorem 4**

Let $(\Theta^{*(k)}, Z^{*(k)}, V^{*(k)})$ be the solution to problem and suppose that $V^{*(k)}$ is not a diagonal matrix. Let $\hat{V}^{(k)} = \text{diag}(V^{*(k)})$, and construct $\hat{Z}$ as follows

$$\hat{Z}_{ij}^{(k)} = \begin{cases} Z_{ij}^{*(k)} + V_{ij}^{*(k)} + V_{ji}^{*(k)} & \text{if} \quad i \neq j \\ Z_{ij}^{*(k)} & \text{otherwise} \end{cases}$$

Then we have $\Theta^{*(k)} = \hat{Z}^{(k)} + \hat{V}^{(k)} + t(\hat{V}^{(k)})$. Thus $(\Theta^{*(k)}, \hat{Z}^{(k)}, \hat{V}^{(k)})$ is also a feasible solution. Now we prove that if the condition holds, $(\Theta^{*(k)}, \hat{Z}^{(k)}, \hat{V}^{(k)})$ has a smaller objective than $(\Theta^{*(k)}, Z^{*(k)}, V^{*(k)})$. We only need to prove that

$$\begin{aligned}
&\lambda_1 \sum_{k=1}^{k} ||\hat{Z}^{(k)} - \text{diag}(\hat{Z}^{(k)})||_1 \\
&+ \lambda_2 \sum_{k<k'} ||\hat{Z}^{(k)} - \hat{Z}^{(k')} - \text{diag}(\hat{Z}^{(k)} - \hat{Z}^{(k')})||_1 \\
\leq\ &\lambda_1 \sum_{k=1}^{k} ||Z^{*(k)} - \text{diag}(Z^{*(k)})||_1 + \\
&+ \lambda_2 \sum_{k<k'} ||Z^{*(k)} - Z^{*(k')} - \text{diag}(Z^{*(k)} - Z^{*(k')})||_1 \\
&+ \lambda_3 \sum_{k=1}^{K} ||V^{*(k)} - \text{diag}(V^{*(k)})||_1 + \lambda_4 \sum_{k=1}^{K} ||V^{*(k)} - \text{diag}(V^{*(k)})||_{1,2} \\
&+ \lambda_5 \sum_{k<k'} ||V^{*(k)} - V^{*(k')} - \text{diag}(V^{*(k)} - V^{*(k')})||_1.
\end{aligned} \tag{21}$$

Two terms in the left hand of formula (21) can be restricted separately.

$$\begin{aligned}
&\lambda_1 \sum_{k=1}^{k} ||\hat{Z}^{(k)} - \text{diag}(\hat{Z}^{(k)})||_1 \\
\leq\ &\lambda_1 \sum_{k=1}^{k} ||Z^{*(k)} - diag(Z^{*(k)})||_1 + 2\lambda_1 \sum_{k=1}^{K} ||V^{*(k)} - diag(V^{*(k)})||_1.
\end{aligned}$$

$$\begin{aligned}
&\lambda_2 \sum_{k<k'} ||\hat{Z}^{(k)} - \hat{Z}^{(k')} - \text{diag}(\hat{Z}^{(k)} - \hat{Z}^{(k')})||_1 \\
\leq \quad &\lambda_2 \sum_{k<k'} ||Z^{*(k)} - Z^{*(k')} - diag(Z^{*(k)} - Z^{*(k')})||_1 \\
&+ 2\lambda_2 \sum_{k<k'} ||V^{*(k)} - V^{*(k')} - diag(V^{*(k)} - V^{*(k')})||_1.
\end{aligned}$$

And

$$\sum_{k=1}^{K} ||V^{*(k)} - diag(V^{*(k)})||_1 \leq \sqrt{p} \sum_{k=1}^{K} ||V^{*(k)} - \text{diag}(V^{*(k)})||_{1,2},$$

by Cauchy inequality.

Then the inequality (21) follows from the sufficient condition.

### S6 Text: Methods for generating the basis proportion for each feature

We generate count data according to two steps. (1) In the first step, we sample basis proportions for each OTU given mean basis abundance and basis covariance, from one of three underlying distributions, such as log ratio normal (LRN), Poisson log normal (LNP), and Dirichlet log normal (LND). (2) For each sample, counts are drawn from Multinomial distribution using the basis proportions obtained in the first step with a sample size. Suppose that basis abundance $a = (a_1, a_2, \ldots, a_p)^T$ has basis covariance $\Sigma_{p*p}$. For L-RN, we sample $\phi_i = \log\left(\frac{x_i}{x_p}\right) \sim MVN(\mu, \Omega)$ for $i = 1, 2, \ldots, p-1$, where $\mu = \log\left(\frac{(a_1, a_2, \ldots, a_{p-1})^T}{a_p}\right), \Omega = L\Sigma L^T$,and

$$L = \begin{pmatrix} 1 & \cdots & 0 & -1 \\ \vdots & \ddots & \vdots & \vdots \\ 0 & \cdots & 1 & -1 \end{pmatrix}_{(p-1)*p}$$

Then, we can obtain basis proportions $w_i = \frac{exp(\phi_i)}{1+\sum\limits_{i=1}^{p-1} exp(\phi_i)}$ for $i = 1, \ldots, p-1$ and $w_p = \frac{1}{1+\sum\limits_{i=1}^{p-1} exp(\phi_i)}$. For LNP, we sample $\phi_i$ such that $\log(\phi_i) \sim MVN(0, \Sigma)$. Then we obtain for each OTU basis abundance $c_i|\phi_i \sim Poisson(a_i\phi_i)$ and proportions $w_i = \frac{c_i}{\sum\limits_{i=1}^{p} c_i}$. For LND, we sample $\phi_i$ such that $\log(\phi_i) \sim MVN(0, \Sigma)$. Then for each sample we draw the basis proportions $w_1, \ldots, w_p|\phi \sim \frac{\Gamma(\sum\limits_{i=1}^{p} \phi_i)}{\prod\limits_{i=1}^{p} \Gamma(\phi_i)} \prod\limits_{i=1}^{p} w_i^{\phi_i - 1}$.

In the second step, the count data are drawn from $x_1, x_2, \ldots, x_p \sim C_X^x \prod\limits_{i=1}^{p} w_i^{x_i}$ for each sample given a sequencing size of $X = \sum\limits_{i=1}^{p} x_i$.

**S7 Text: A simple example computation of TPR and FPR based on current method**

Below we explain how we compute the TPR (True Positive Rate), FPR (False Positive Rate) and Precision with an example.

Suppose the nodes are labelled as $1, 2., ..., p$ and our program detects the following nodes as hubs under three conditions [1],[2],[3]:

```
[1] 7, 15, 22, 36, 60, 77, 79, 84, 145, 154
[2] 15, 22, 60, 77, 79, 98, 145, 154
[3] 12, 15, 22, 60, 67, 100, 109, 135, 145, 154, 159
```

Suppose the true hubs are located at these nodes under three conditions [1],[2],[3]:

```
[1] 7, 15, 22, 36, 60, 79, 84, 154
[2] 15, 36, 60, 77, 79, 98, 145, 154
[3] 15, 22, 60, 100, 109, 145, 154, 159
```

When the relevant elements to detect are the (true) common hubs 15, 60, 154, our program successfully detects node 15, 60, 154, so

$$\text{TPR-C} = \frac{TP}{P} = 1$$

Out of the $p - 3$ node locations which are not true common hubs, our program falsely detects node 22, 145 as common hubs, so

$$\text{FPR-C} = \frac{FP}{N} = \frac{2}{p-3}$$

$$\text{Precision-C} = \frac{TP}{TP + FP} = \frac{3}{3+2}$$

When the relevant elements to detect are the eleven (true) conditional-specific hubs 36,79 (under [1,2]), 145 (under [2,3]), and 22 (under [1,3]), 7, 84 (under [1]), 22 (under [2]), 100, 109, 159 (under [3]), our program successfully detects nodes 7, 79, 84, 98, 100, 109, 159 so

$$\text{TPR-S} = \frac{TP}{P} = \frac{7}{11}$$

Out of the $p - 11$ nodes which are not conditional-specific hubs (including all nodes which are either true non-hubs or true common hubs), our program makes one mistake: it falsely detects true common hub 12, 67, 135 as condition [3]-specific. So

$$\text{FPR-S} = \frac{FP}{N} = \frac{3}{p-11}$$

$$\text{Precision-S} = \frac{TP}{TP+FP} = \frac{7}{7+3}.$$

Overall, the number of true hubs for each class is eight, and we successfully detect 8, 7 (missing 36), 8 respectively. Out of the $3p - 24$ nodes we falsely identify 77, 145 (under [1]), 22 (under [2]), 12, 67, 135 (under [3]). The corresponding total TPR, FPR and Precision are compute as

$$\text{TPR} = \frac{8+7+8}{8+8+8}$$

$$\text{FPR} = \frac{2+1+3}{3p-24}$$

$$\text{Precision} = \frac{8+7+8}{10+8+11}.$$

**S1 Table: Additional simulations for only common hubs and only class-specific hubs**

Table 1: The ability of EDOHA, JRmGRN and HGL to identify hub nodes in the networks with "only common hubs" are shown by True Positive (TP), False Positive (FP), sensitivity and specificity. We generate 100 datasets with sample size n=80.

| | 80 nodes: 5 hubs | | 160 nodes: 8 hubs | | 300 nodes: 12 hubs | |
|---|---|---|---|---|---|---|
| | TPR | FPR | TPR | FPR | TPR | FPR |
| EDOHA | 0.821 | 0.013 | 0.796 | 0.003 | 0.763 | 0.001 |
| JRmGRN | 0.895 | 0.017 | 0.917 | 0.016 | 0.875 | 0.004 |
| HGL | 0.753 | 0.024 | 0.769 | 0.002 | 0.662 | 0.001 |

Table 2: The ability of EDOHA and JRmGRN to identify hub nodes in the networks with “only class-specific hubs” are shown by True Positive (TP), False Positive (FP), sensitivity and specificity. We generate 100 datasets with sample size n=80.

| | 80 nodes: 5 hubs | | 160 nodes: 8 hubs | | 300 nodes: 12 hubs | |
|---|---|---|---|---|---|---|
| | TPR | FPR | TPR | FPR | TPR | FPR |
| EDOHA | 0.871 | 0.027 | 0.746 | 0.026 | 0.711 | 0.025 |
| HGL | 0.879 | 0.046 | 0.797 | 0.021 | 0.715 | 0.01 |
| JRmGRN | NA | NA | NA | NA | NA | NA |

**S1 Fig: Venn diagram of consistent correlated OTUs from Control, Healthy and EBA groups**

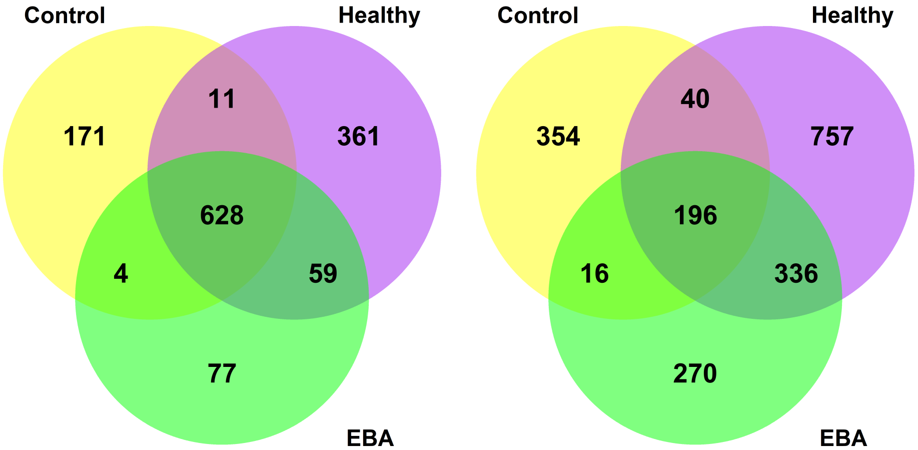


(a) Our results (b) Ban et al. results

**S2 Table: The hub proteins detected in four organs and their functions in living organism**

| Hub | Tissue | Function |
|---|---|---|
| DDX21 | Colon; Liver; Lung; Kidney | play an important role in ribosomal RNA biogenesis, RNA editing, RNA transport, and general transcription |
| REEP6 | Colon; Liver; Lung; Kidney | the transport of receptors from the endoplasmic reticulum (ER) to the cell surface and regulates ER membrane structure |
| SEPSECS | Colon; Liver; Lung; Kidney | convert O-phosphoseryl-tRNA(Sec) to selenocysteinyl-tRNA(Sec) required for selenoprotein biosynthesis |
| TIMM9 | Liver; Lung; Kidney | mediate the import and insertion of hydrophobic membrane proteins into the mitochondrial inner membrane |
| HMOX1 | Liver; Kidney | an essential enzyme in heme catabolism |
| PRKAR2B | Colon; Liver | regulatory subunit of the cAMP-dependent protein kinases involved in cAMP signaling in cells, and cAMP is a signaling molecule important for a variety of cellular functions |
| MRPS5 | Colon | RNA binding and structural constituent of ribosome |
| BCKDK | Liver | the key regulatory enzyme of the valine, leucine and isoleucine catabolic pathways and regulates the activity state of the BCKD complex |
| COMT | Liver | participate in the metabolism of endogenous substances |
| BZW2 | Lung | be involved in cell differentiation and nervous system development |
| SLC44A2 | Lung | exhibit some choline transporter activity |
| STOM | Lung | regulate ion channel activity and transmembrane ion transport |

| | | |
|---|---|---|
| ATP1B1 | Kidney | the non-catalytic component of the active enzyme, which catalyzes the hydrolysis of ATP coupled with the exchange of $Na^+$ and $K^+$ ions across the plasma membrane |
| ATP6AP1 | Kidney | be required for luminal acidification of secretory vesicles. |
| ATP6V1A | Kidney | mediate acidification of eukaryotic intracellular organelles and play a part in neurite development and synaptic connectivity |
| CCDC86 | Kidney | be involved in RNA binding and viral process |
| ETFA | Kidney | be required for normal mitochondrial fatty acid oxidation snd normal amino acid metabolism |
| NUP210 | Kidney | a membrane-spanning glycoprotein that is a major component of the nuclear pore complex |
| PTGES2 | Kidney | Isomerase that catalyzes the conversion of PGH2 into the more stable prostaglandin E2 (PGE2) |
| SCARB1 | Kidney | a plasma membrane receptor for high density lipoprotein cholesterol (HDL) mediate cholesterol transfer to and from HDL |

**Reference**


[1] Stephen Boyd, Stephen P Boyd, and Lieven Vandenberghe. *Convex optimization.* Cambridge university press, 2004.

[2] Patrick Danaher, Pei Wang, and Daniela M Witten. The joint graphical lasso for inverse covariance estimation across multiple classes. *Journal of the Royal Statistical Society: Series B (Statistical Methodology)*, 76(2):373–397, 2014.

[3] Holger Hoefling. A path algorithm for the fused lasso signal approximator. *Journal of Computational and Graphical Statistics*, 19(4):984–1006, 2010.
